# Switching from tonic to burst firing creates an attractor in synaptic weight dynamics

**DOI:** 10.1101/2022.07.15.500198

**Authors:** Kathleen Jacquerie, Danil Tyulmankov, Pierre Sacré, Guillaume Drion

## Abstract

1

Neural circuits often alternate between tonic and burst firing – two distinct activity regimes that reflect changes in excitability and neuromodulatory state. While tonic firing produces asynchronous spiking driven by diverse external inputs, collective burst firing consists of rapid clusters of spikes followed by a period of silence, happening synchronously within the network. Synaptic plasticity has typically been studied only in either one of these regimes, leaving unclear how alternating states jointly shape long-term weight dynamics. Here, we use a conductance-based network model endowed with calcium-based or spike-timing–based plasticity rules to examine how synaptic weights evolve across state transitions. During tonic firing, synaptic weights are driven by the statistics of external inputs, producing a broad distribution across the network. In contrast, during collective burst firing, weights converge to a narrow region in weight space: a burst-induced attractor. We derive the location of this attractor analytically in terms of plasticity parameters and activity statistics, and confirm its emergence across diverse plasticity rules. The attractor reflects the synchronization of plasticity-driving signals during bursts, which homogenizes synaptic dynamics and forces convergence toward shared fixed points. We further show that neuromodulation and synaptic tagging can shift or split the burst-induced attractor, stabilizing selected synapses while weakening others. This mechanism reconciles flexibility during tonic-driven learning with stability during burst-driven consolidation. These results identify burst-induced attractors as a robust emergent property of networks combining collective bursting with soft-bound plasticity rules. By showing how they can be analytically predicted and experimentally modulated, our work provides a general computational framework linking state transitions, synaptic plasticity, and memory organization.

**Author Summary:** Brains operate in different activity states, reflecting different behaviors or neuromodulatory states. Neurons can fire isolated spikes in a tonic mode that encodes information about external inputs. They can fire rapid bursts of spikes, generating large synchronized oscillations that dominate population activity. Both tonic and burst firing are linked to learning and memory, yet their distinct contributions to shaping synaptic plasticity remain poorly understood. In this study, we use biophysical network models equipped with well-established plasticity rules to investigate how synaptic weights evolve under tonic and burst firing. We show that during tonic activity, synapses diverge toward a wide variety of values, reflecting the diversity of input statistics. In contrast, when the network enters a collective bursting state, synaptic weights collapse into a narrow region of weight space—a “burst-induced attractor.” We derive the attractor mathematically and show that its position depends directly on the plasticity parameters, meaning it can be shifted or split through neuromodulatory and tag-dependent processes. Our results suggest that bursts provide a robust and controllable stage for synaptic consolidation.

**Graphical abstract:** 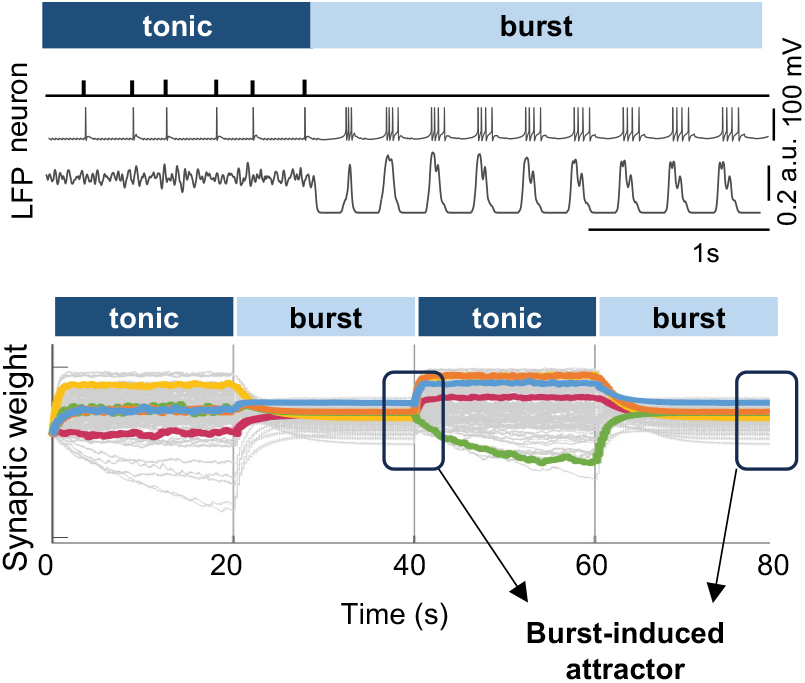

## 3 Introduction

Neural circuits can alternate between distinct neuronal firing regimes—most notably either tonic or burst firing—depending on behavioral demands and neuromodulatory state (McGinley et al., 2015). In tonic firing, neurons emit relatively isolated spikes that are primarily driven by ongoing external inputs. At the population level, this activity corresponds to low-amplitude, high-frequency oscillations in local field potential (LFP) or electroencephalogram (EEG) recordings. In contrast, burst firing consists of rapid clusters of spikes followed by periods of silence. Bursts can emerge endogenously from intrinsic neuronal and network properties, reflecting cycles of ion channel activation and inactivation, like T-type calcium channels (McCormick and Salkoff, 2015; Lee and Dan, 2012; Dastgheib et al., 2022; Jacquerie and Drion, 2021). When many neurons burst synchronously, the resulting collective dynamics generate large-amplitude, low-frequency LFP/EEG signals characteristic of highly synchronized activity. This dynamic switching between tonic and burst regimes is observed across multiple neural systems and is thought to support different functional states of the brain. Tonic firing is typically associated with active sensory processing and learning (Sherman, 2001), whereas burst firing is more often observed during quiet behavioral states that facilitate memory consolidation (McGinley et al., 2015; Timofeev et al., 2002).

Neural circuits store information via synaptic plasticity—activity-dependent changes in synaptic strength. These changes involve multiple mechanisms, including increased postsynaptic receptor efficacy, the insertion of additional receptors via exocytosis, de novo protein synthesis, and structural remodeling of synapses (Lamprecht and LeDoux, 2004; Fauth and Tetzlaff, 2016; Citri and Malenka, 2008; Magee and Grienberger, 2020). Synaptic plasticity, however, has largely been studied under either tonic or burst firing in isolation, and it remains unclear whether these regimes drive different forms of information encoding or learning; whether tonic and burst firing support distinct modes of synaptic weight change, or whether they engage overlapping mechanisms. Our central question is therefore: how do synaptic weights evolve when activity shifts between tonic and burst firing, and what implications does this have for learning and memory?

This question has largely been unexplored in computational models, primarily for two reasons. First, it requires a network model capable of robustly alternating between firing modes despite variability in neuronal and synaptic properties. Second, it requires a plasticity rule that remains valid across distinct activity regimes. Most existing models address plasticity in only one regime at a time (Tyulmankov, 2025). Many rely on spike-timing–dependent plasticity (STDP) formulations—pair-based or triplet rules (Song et al., 2000; Van Rossum et al., 2000; Pfister and Gerstner, 2006)—or on calcium-based plasticity models (Shouval et al., 2002; Graupner et al., 2016; Deperrois and Graupner, 2020), which are typically tested under tonic firing conditions. Burst-related plasticity has also been explored using burst-timing–dependent rules (Gjorgjieva et al., 2009). However, neither approach has been examined in the context of networks switching between firing states, despite the prevalence of such state transitions in the brain.

To address this gap, we build on our previous work, which introduced a biophysical, conductance-based network that robustly switches between tonic and burst firing (Jacquerie and Drion, 2021). Here we endow that network with synaptic plasticity using a calcium-based rule (Graupner et al., 2016; Graupner and Brunel, 2012) and examine, in parallel, representative spike-based rules (pair-based and triplet STDP).

We find that when the network enters a collective bursting state, synaptic weights converge to a narrow region in weight space—a burst-induced attractor. The same qualitative phenomenon emerges across calcium- and STDP-inspired rules, and we derive the attractor location analytically. Crucially, the attractor follows directly from the plasticity parameters, so it can be modified by neuromodulatory and tag-dependent processes. This turns the attractor into a feature: during bursting, adjusting plasticity parameters globally or in a tag-dependent manner can shift or split the attractor, enabling selective consolidation (e.g., stabilizing tagged synapses while down-selecting untagged ones). More broadly, this frames bursting as a plausible controllable consolidation stage.

Overall, our results argue that plasticity should be studied in networks that alternate between tonic and burst firing, and they provide a general mechanism linking collective burst firing to synaptic organization and consolidation.

## 4 Results

### 4.1 Modeling robust transitions between tonic and burst firing in a biophysical network model

We simulate the transition from tonic to burst firing in a heterogeneous network of conductance-based model neurons (Sections 6.1.1, 6.1.4), with the network architecture shown in Fig 1A. The network is composed of *N* presynaptic excitatory neurons projecting to *M* postsynaptic excitatory neurons with membrane potentials *V*_*j*_ and *V*_*i*_, respectively. These neurons all receive GABA currents from a single inhibitory neuron (*V*_inh_), the activity of which controls whether the network is in a tonic firing or bursting regime. This modulation in inhibitory levels mimics the variation of neuromodulators such as acetylcholine, dopamine, serotonin, or histamine (Tyree and Luis De Lecea, 2017), known to control brain state transitions (Hill and Tononi, 2005; Bazhenov et al., 2002; Olcese et al., 2010; Jacquerie and Drion, 2021; Drion et al., 2018). Each neuron also receives an individual applied current (*I*_app,inh_, *I*_app,*j*_, *I*_app,*i*_) to set its firing rate for learning specified input patterns. For details on equations goverining neural dynamics, see Methods section 6.1.3.

**Fig. 1:**
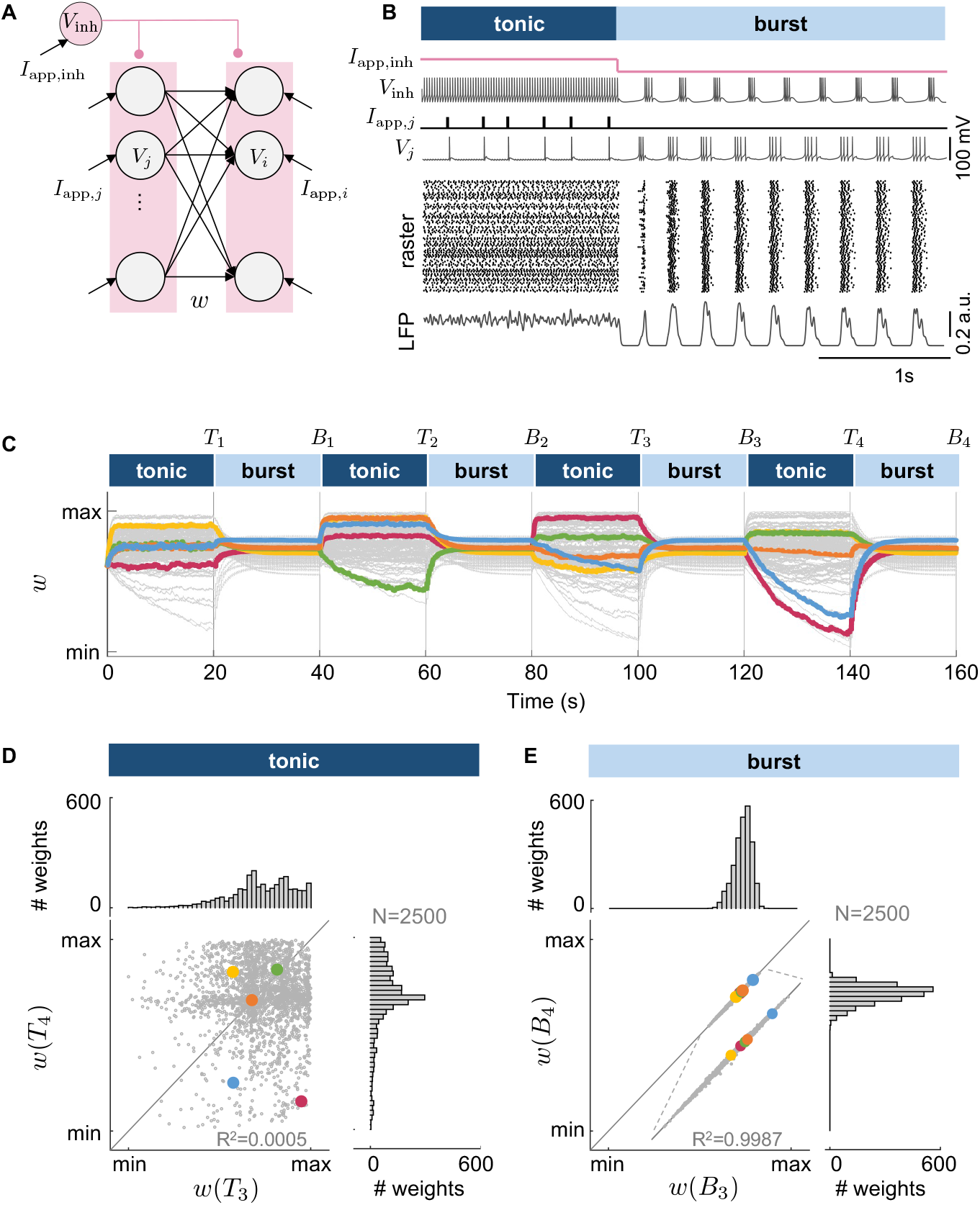
Switching from tonic to burst drives the synaptic weights towards an attractor in the weight space. **A**. The network model consists of one inhibitory neuron (*V*_inh_) projecting to all the excitatory neurons, connected in a feedforward configuration with *N* presynaptic neurons (*V*_*j*_) to *M* postsynaptic neurons (*V*_*i*_). An external current applied to the inhibitory neuron controls the network state (*I*_app,inh_). Each excitatory neuron is subjected to an individual external current mimicking an external input. **B**. Hyperpolarizing the current applied to the inhibitory neuron (*I*_app,inh_) switches the network state from input-driving tonic firing to collective burst firing. Raster plot activity of the excitatory neurons and local field potential (LFP) traces are displayed. **C**. Evolution of the synaptic weights during switches from input-driven tonic firing to collective burst firing. The network comprises 50 presynaptic neurons connected to 50 postsynaptic neurons for 100 traces among 2500 (gray lines) with 5 randomly highlighted lines (colored lines). *T*_*k*_ and *B*_*k*_ indicate the end of the *k*-th tonic and burst firing state, respectively. **D-E**. Comparison of the synaptic weights at the end of the third and fourth tonic firing states (dark blue) or the third and fourth burst firing states (light blue), normalized between the minimal and maximal values shown on a scatter plot. The histograms highlight the weight distribution. The weights converge to a fixed point in the weight space, we call burst-induced attractor (min(*w*_*ij*_) = 0.176, max(*w*_*ij*_) = 0.705).

Fig 1B illustrates the transition between the two firing regimes in a network of 50 presynaptic and 50 postsynaptic neurons: *input-driven tonic firing* and *collective burst firing*, controlled by the current *I*_app,inh_ applied to the inhibitory neuron.

During *input-driven tonic firing*, a depolarizing current *I*_app,inh_ applied to the inhibitory neuron induces regular tonic spiking activity, as commonly observed in inhibitory pacemaking cortical neurons (McGinley et al., 2015). Consequently, this pacemaking inhibitory neuron sets the resting membrane voltage of all the excitatory neurons. Upon receiving an applied current (*I*_app,*j*_, *I*_app,*i*_), these excitatory neurons generate action potentials. For details on protocol inducing tonic firing, see Methods section 6.1.2).

To trigger a network transition from input-driven tonic firing to *collective burst firing*, a hyperpolarizing current *I*_app,inh_ is applied to the inhibitory neuron (Section 6.1.2). This bursting activity—in which each neuron generates a rapid succession of action potentials followed by a quiescent period (Zeldenrust et al., 2018; Desroches et al., 2022; Fernandez and Luthi, 2019)—is independent of external input currents *I*_app,*j*_ and *I*_app,*i*_, and is caused by neuron intrinsic properties, such as T-type calcium channels that deinactivate when the neuron is hyperpolarized. The term “collective” refers to the synchronization of individual bursting neurons despite heterogeneity in intrinsic properties and synaptic connections, resulting in slow large-amplitude population activity shown in the LFP oscillation, contrasting the fast small-amplitude observed in input-driven tonic firing (Fig 1B, LFP traces) (Jacquerie and Drion, 2021). For details on LFP calculation, see Methods section 6.1.5.

### 4.2 Modeling synaptic plasticity

Synapses between presynaptic and postsynaptic excitatory neurons undergo activity-dependent plasticity— a fundamental process underlying learning and memory. In biological systems, plasticity arises from a combination of biochemical, structural, and electrical mechanisms that respond to intracellular calcium dynamics.

To capture these dynamics quantitatively, we use a state-of-the-art calcium-based model established by Graupner et al. (Graupner and Brunel, 2012), which was fit to experimental data obtained through a frequency pairing protocol (Sjöström et al., 2001). The model implements two opposing calcium-triggered pathways that lead to either an increase or a decrease in synaptic strength: potentiation or depression is triggered when calcium levels exceed specific thresholds. Although we illustrate our results with this model, we later show that the same principles apply across a wide range of plasticity rules, highlighting the generality of our results. In this framework, the synaptic weight from the presynaptic neuron *j* to the postsynaptic neuron *i* is denoted *w*_*ij*_ and its evolution is governed by Equation 3. For details on equations governing synaptic plasticity, see Methods section 6.1.6.

### 4.3 Collective burst firing drives synaptic weights towards an attractor in the weight space

To study the interaction between switches in firing activity and synaptic plasticity, we track the evolution of the weights over the course of eight transitions between tonic and burst firing (Fig 1C).

During tonic firing (Fig 1C, dark blue), each neuron receives a random pulse train input, with the stimulation frequency selected uniformly at random between 0.1 and 50 Hz. For details on the computational experiment, see Methods section 6.1.7. The temporal evolution of the synaptic weight *w*_*ij*_(*t*) is determined by the correlation between the activity of presynaptic neuron *j* and postsynaptic neuron *i*, which drives plasticity at those synapses. As a result, at the end of each tonic firing period (time *T*_*k*_, *k* = 1, …, 4), the weight *w*_*ij*_(*T*_*k*_) converges to a value determined by the specific external input received. Since different neurons receive different input statistics, the weights across the network settle at a broad range of values (see Fig 1D, histograms). Furthermore, because the input statistics vary across tonic periods, the synaptic weights also differ from one tonic episode to the next, as illustrated by plotting *w*(*T*_4_) against *w*(*T*_3_) (Fig 1D, scatter plot; see also Fig S1 for comparisons across other states).

During burst firing (Fig 1C, light blue), the network is driven only by an inhibitory current, with no external input to the excitatory neurons. Collective bursting activity drastically alters synaptic weight dynamics: previously potentiated connections tend to depress, and vice versa. Each initial weight *w*_*ij*_(*T*_*k*_) converges to a unique steady-state value *w*_*ij*_(*B*_*k*_) within a narrow range (Fig 1E, histograms), and repeats from one burst episode to the next one independent of its initial condition, as illustrated by plotting *w*(*B*_4_) against *w*(*B*_3_)(Fig 1E, scatter plot; see Fig S1 for other pairs). We call this phenomenon a *burst-induced attractor* and define it as a narrow region in weight space to which synaptic weights converge under collective bursting dynamics.

### 4.4 The burst-induced attractor emerges in a wide range of synaptic plasticity models

We next investigate whether the choice of synaptic plasticity rule influences the emergence of a burst-induced attractor. We repeat the same experiment in the same network using various synaptic plasticity models and observe that the effect persists across a wide range of rules (Fig S2), including different variants of calcium-based models (Graupner et al., 2016; Shouval et al., 2002; Deperrois and Graupner, 2020), phenomenological models based on pairwise (Song et al., 2000) or triplet (Pfister and Gerstner, 2006) spike timing, weight-dependent rules (Van Rossum and Shippi, 2013; Gütig et al., 2003), and models with parameters fitted to cortical frequency-based protocols (Sjöström et al., 2001) or hippocampal spike-based protocols (Bi and Poo, 1998). We also compare the influence of weight dependence by testing both soft and hard bounds in the synaptic rule. During collective burst firing, synaptic weights converge towards an attractor in the weight space no matter the choice of the synaptic plasticity rule.

### 4.5 Analytical insight into the burst-induced attractor

Why does collective burst firing constrain synaptic weights into an attractor, whereas tonic firing produces a broad spread? To address this question, we derive the fixed points of synaptic weight dynamics under calcium-based and spike-timing–based plasticity rules and then verify these predictions in network simulations.

We first focus on calcium-based plasticity rules, and we derive the fixed point for one of the rules, (Graupner et al., 2016). Two thresholds determine the outcome of calcium dynamics: a depression threshold *θ*_d_ and a potentiation threshold *θ*_p_. Each calcium regime is associated with a steady-state value (potentiation or depression) and a characteristic relaxation timescale. When calcium remains below *θ*_d_ (*c*_*ij*_ *< θ*_d_), the weight does not change. For intermediate calcium levels (*θ*_d_ ≤ *c*_*ij*_ *< θ*_p_), the weight drifts toward the depression steady state Ω_d_ with time constant *τ*_d_. When calcium exceeds *θ*_p_ (*c*_*ij*_ ≥ *θ*_p_), the weight relaxes toward the potentiation steady state Ω_p_ with time constant *τ*_p_. As shown in Fig 2A, calcium fluctuations can therefore be decomposed into potentiation, depression, and neutral intervals. By averaging the fast calcium dynamics, we obtain the fixed point of the synaptic weight:

**Fig. 2:**
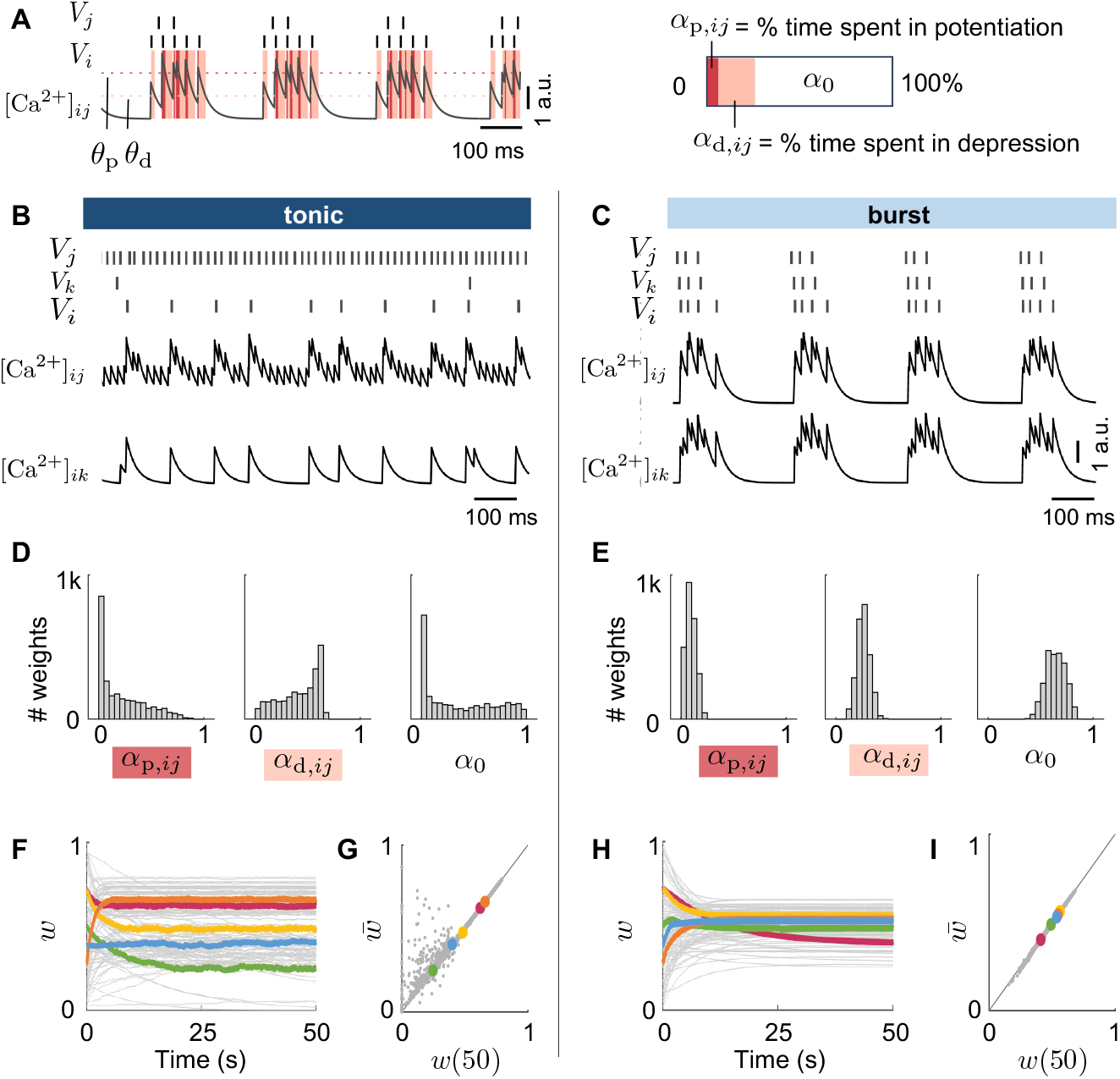
Burst firing constrains synaptic weights toward an attractor via shared calcium dynamics. **A.A**. Calcium trace decomposition into potentiation, depression, and neutral zones, based on pre- and postsynaptic spike timing. Red and orange shading indicate time spent above the potentiation threshold or below the depression threshold, respectively. Right: effective time in each region, weighted by corresponding time constants (*α*_p,*ij*_, *α*_d,*ij*_). **B-C**. Example spike trains and calcium traces for two synapses during tonic and burst firing. Tonic activity produces diverse calcium dynamics, while burst firing induces synchronized fluctuations across synapses. **D-E**. Distribution of effective time allocated in different plasticity regions during tonic and burst firing. Tonic firing shows broad distributions, while burst firing produces narrower, more consistent distributions across synapses. **F-H**. Evolution of synaptic weights over time during tonic (F) and burst (H) firing, starting from the same initial conditions. Weights diverge toward distinct values under tonic input, whereas they converge to a narrow range under burst firing, indicating the emergence of a burst-induced attractor. **G-I**. Predicted steady-state weight using analytical equations (Equation 1) compared with actual simulated weights at different time steps. The linear regression between the predicted values and the actual final values at the end of the stimulation is closer in burst firing compared to tonic firing (*R*^2^ = 0.9983 in burst and *R*^2^ = 0.9100 in tonic).

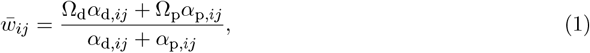

where we define the effective times spent in the depression and potentiation regimes (*α*_d,*ij*_ and *α*_p,*ij*_, respectively). This derivation is based on previous works (Dorman and Blackwell, 2021; Gütig et al., 2003). For more details on the derivation, see section Text S4.

A similar principle holds for STDP rule, where correlations between pre- and postsynaptic spikes reflect how much time is spent in potentiation versus depression window. For the STDP model of (Van Rossum et al., 2000), the fixed point is:

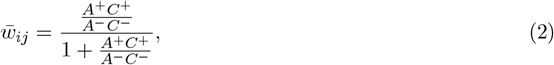

where *A*^+^ and *A*^−^ are potentiation and depression parameters of the STDP rule, and *C*^+^ and *C*^−^ are measures of the synaptic potentiation and depression that stems from all input–output correlations with positive and negative time lag, respectively. For more details on the derivation, see section Text S5. The results for hard-bounds can be found in section Text S6.

In both cases, the fixed point depends on the plasticity parameters and on activity statistics—time spent in calcium regimes or spike correlations (Gütig et al., 2003; Jedrzejewska-Szmek et al., 2017).

We test these predictions in the generic network shown in Fig 1A, consisting of 50 presynaptic and postsynaptic neurons. The network operates either in tonic mode, with random inputs (0.1–50 Hz), or in burst mode, induced by hyperpolarizing the inhibitory neuron. For details on the computational experiment, see Methods section 6.1.8.

During tonic firing, inputs are uncorrelated, so each synapse experiences its own calcium pattern. As shown in Fig 2B-C, calcium traces differ widely across synapses. The resulting effective times in potentiation and depression are broadly distributed (Fig 2D), and weights converge to a wide range of fixed points (Fig 2F). In contrast, burst firing synchronizes activity, producing similar calcium traces and correlations across the network (Fig 2E). Consequently, all synapses converge toward a narrow set of fixed points—the burst-induced attractor (Fig 2H).

Analytical predictions closely match simulated weights (Fig 2G–I), with stronger correspondence in bursts (*R*^2^ = 0.9983) compared to tonic activity (*R*^2^ = 0.9100). The lower *R*^2^ in tonic reflects finite-time effects: some synapses are predicted to depress toward zero but have not yet completed that trajectory within the 50 s window, leading to larger residuals. The STDP model shows the same pattern—lower in tonic (*R*^2^ = 0.3104) and high in burst (*R*^2^ = 0.9747)—for the same reasons and further underscores that an attractor yields a clear, accurate prediction in the burst regime.

Together, these results show that burst firing creates an attractor in weight space because shared activity patterns synchronize the effective plasticity drive across synapses, forcing them toward similar fixed points. Tonic firing, by contrast, disperses weights because each synapse follows its own independent activity statistics.

### 4.6 Burst-induced attractor sensitivity to rhythmic state modulation

We next apply our analytical result to show how the burst-induced attractor can be modulated in practice, turning the apparent “reset” into a flexible mechanism for consolidation. Our framework predicts that altering the collective bursting dynamics modifies the effective time spent in potentiation and depression regimes (in calcium-based rules) or the spike-time correlations (in spike-based rules). As a result, the location of the attractor can be shifted without any other alteration.

To illustrate this, we return to the network described in Fig 1A. Then, we emulate changes in the neuromodulatory drive by applying three different levels of external current to the inhibitory neuron (Fig 3A, *I*_app,inh_). This manipulation mimics changes in neuromodulatory drive and produces distinct patterns of collective bursting activity, as seen in the raster plots and LFP profiles (Fig 3A). For details on the computational experiment, see Methods section 6.1.9. Consistent with the analytical prediction, the attractor shifts slightly depending on the bursting regime, yet synaptic weights still converge within a narrow region of weight space (Fig 3A, bottom). By contrast, synaptic weights at the end of tonic firing remain broadly distributed, highlighting that bursting—despite resetting weights—can reorganize them in a structured and controllable way (Fig 3B).

**Fig. 3:**
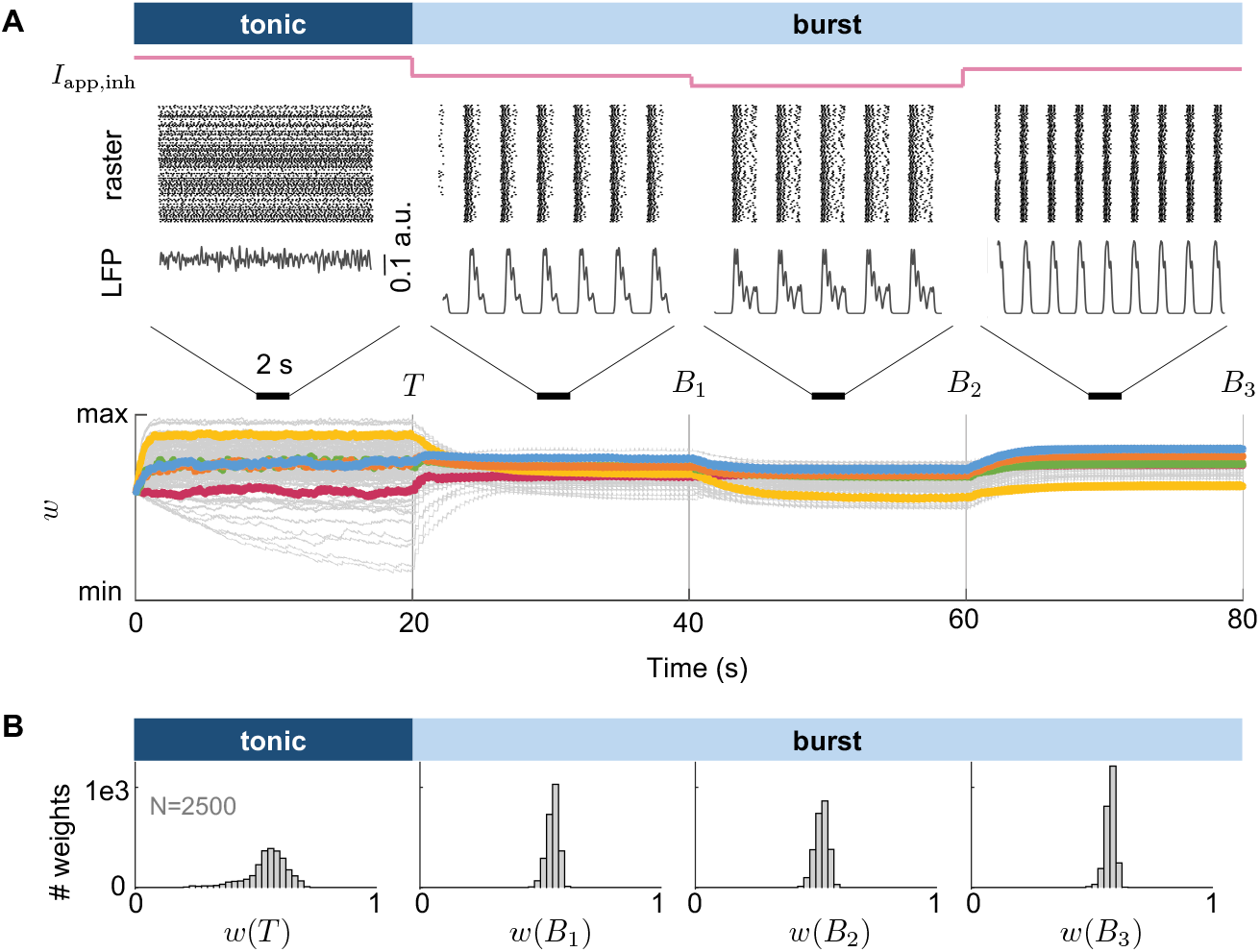
Modulation of burst-induced attractor in synaptic weights. **A**. Evolution of the network activity and synaptic weights when the network transitions from tonic to burst firing with three variations of the collective burst firing (achieved by modulating the external current applied to the inhibitory neuron). T indicates the end of the tonic firing state, and *B*_*k*_ indicates the end of the *k*-th burst firing state. 100 traces are randomly displayed among 2500 synaptic connections. The steady-state values associated can be modulated by the collective burst firing, moving the attractor in the weight space. min(*w*_*ij*_)=0.315 and max(*w*_*ij*_) = 0.703. **B**. Distribution of the synaptic weights at the end of each state.

### 4.7 Neuromodulated and tag-dependent synaptic plasticity rules can split and shift the burst-induced attractor for memory consolidation

We can show the potential of the burst-induced attractor as a mechanism for synaptic consolidation and selection by applying our analytical result (Equation 1 and Equation 2) to show how the burst-induced attractor can be modulated, in practice, by altering the plasticity parameters (e.g. Ω_*p*_ or Ω_*d*_ for calcium-based rule or *A*^*±*^ or *τ*^*±*^ for STDP-like rule).

Recent evidence shows that synaptic plasticity is modulated by neuromodulators such as acetylcholine, dopamine, noradrenaline, serotonin, and histamine (Brzosko et al., 2019; Zannone et al., 2018; Foncelle et al., 2018; Salgado et al., 2012; Nadim and Bucher, 2014; Lisman et al., 2011; Pawlak, 2010; Kirkwood, 2007; Shine et al., 2021; Gerstner et al., 2018; Izhikevich, 2007). These neuromodulators influence plasticity at multiple stages of induction and consolidation (Seol et al., 2007; Bazzari and Parri, 2019; Pawlak, 2010; Nadim and Bucher, 2014). Several computational models have incorporated neuromodulated plasticity rules (Butson, 2012; Frémaux and Gerstner, 2015). For example, in sleep-dependent memory consolidation, the classical STDP kernel observed during wakefulness can be replaced with a purely depressive kernel during sleep, promoting synaptic down-selection (González-Rueda et al., 2018). Other models have shown that acetylcholine or dopamine can dynamically shift the STDP kernel to explain receptive field plasticity and reward-driven navigation (Brzosko et al., 2017; Pedrosa and Clopath, 2017). Additional frameworks introduce neuromodulation via eligibility traces or third-factor models (Gerstner et al., 2018; Izhikevich, 2007; Foncelle et al., 2018).

Therefore, we model a simplified circuit of one inhibitory neuron connected to two excitatory neurons. Two consecutive 30-second periods of tonic and burst firing were simulated. Six circuit instances were tested: three with highly correlated excitatory firing (25–60 Hz) and three with low correlation (≤ 10 Hz). For details on the computational experiment, see Methods section 6.1.10. During the second burst epoch, the potentiation level Ω_p_ was reduced from 0.65 to 0.2, producing a global shift in the burst-induced attractor (Fig 4B). By modulating these parameters, neuromodulators can shift the burst-induced attractor in synaptic weight space. This effect can be replicated in spike-based rules by decreasing the amplitudes of potentiation (*A*^+^) and depression (*A*^−^) in the STDP kernel (Fig S3).

**Fig. 4:**
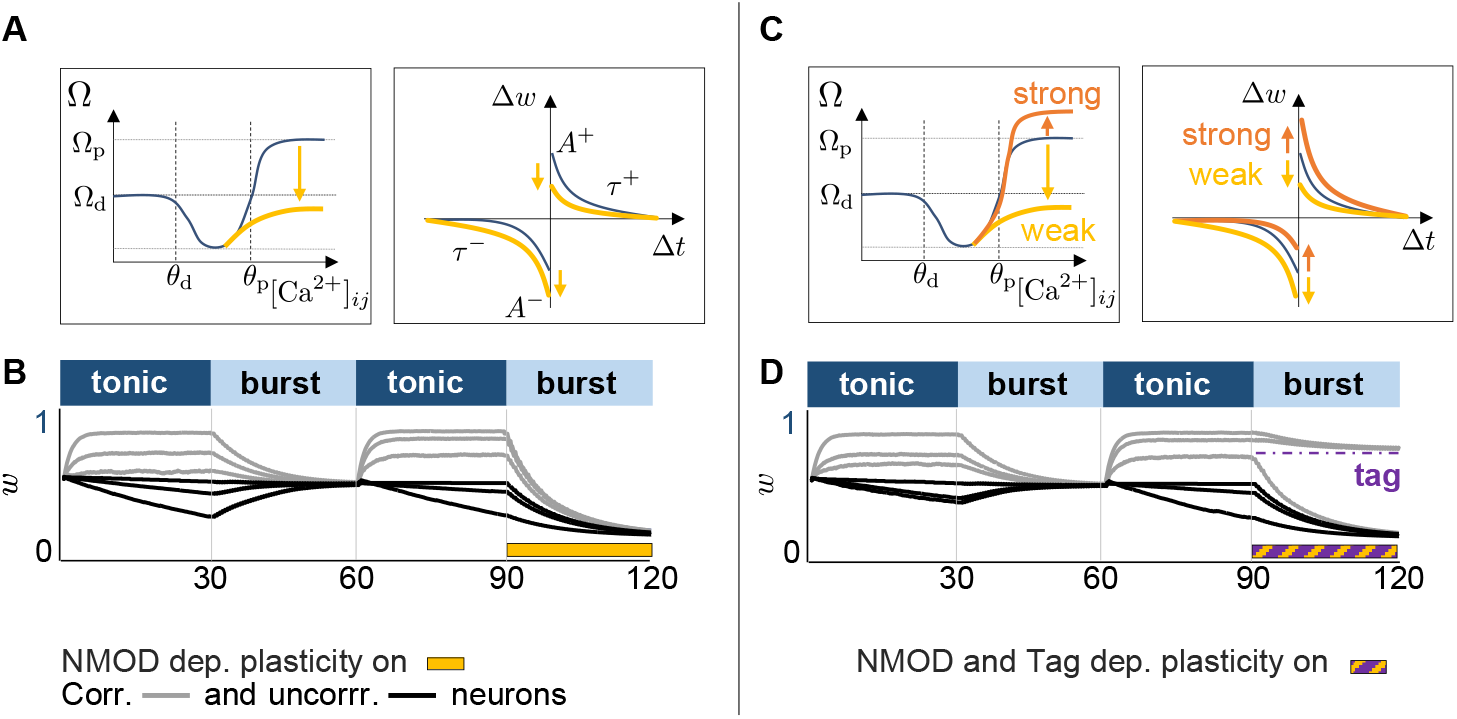
Neuromodulated and tag-dependent synaptic plasticity rules exploit the burst-induced attractor for memory consolidation. **A**. Calcium-based (left) and spike-based (right) plasticity rules, in which potentiation and depression parameters (Ω^p^, *A*^+^, *A*^−^) are modified by neuromodulation. **B**. Evolution of synaptic weights in six circuits undergoing two tonic–burst transitions. In the last burst epoch, the plasticity rule is down-neuromodulated (yellow bar), leading to a different lower burst-induced attractor. Half of the circuits display correlated excitatory activity (gray) and half uncorrelated activity (black). **C**. Calcium-based (left) and spike-based (right) plasticity rules with tag-dependent neuromodulation. Tagged synapses (orange) receive up-neuromodulation, while untagged synapses (yellow) receive down-neuromodulation. **D**. Same protocol as in **B**, but with tag-dependent neuromodulation during the last burst epoch (yellow–purple hatched bar). Synapses tagged during the tonic state (weights above threshold) converge to a higher burst-induced attractor, while untagged synapses converge to a lower one, resulting in bimodal weight consolidation.

We next apply weight-dependent parameter changes only to synapses that exceed a tagging threshold at the end of the preceding tonic phase. Tagged synapse experience enhanced potentiation parameters (Ω_p_ = 1), while untagges synapses are downscaled (Ω_p_ = 0.2).

This selective modulation stabilizes tagged synapses near the higher values attractor and drives untagged synapses toward the lower one, yieleding to a bimodal distribution (Fig 4D). This tag-depdent burst-induced attractor implements selective stabilization for strong synapses while weaking others, a computational realization consistent with synaptic tagging and capture (Frey and Morris, 1997; Redondo and Morris, 2011; Seibt and Frank, 2019; Okuda et al., 2021).

Importantly, this process is not limited to bimodal outcomes. Because the attractor depends directly on the chosen parameters, any mapping from synaptic weight to parameter values can produce different target distributions during bursting. The resulting distribution can therefore be unimodal, bimodal, multimodal, or graded.

Together, these results show that neuromodulation and tagging make the apparent reset during bursting a flexible consolidation stage. Bursting compresses weights into an attractor, but the attractor itself can be shifted or split into several stable states depending on parameter changes.

## 5 Discussion

In this work, we investigate how tonic and burst firing differentially shape synaptic plasticity within a biophysical network model capable of robustly switching between these two activity regimes (Jacquerie and Drion, 2021). We endow this network with a calcium-based plasticity rule (Graupner et al., 2016; Graupner and Brunel, 2012) to explore how each regime contributes to the organization of synaptic weights. We also compare different synaptic plasticity rules such as pair-based or triplet to generalize our outcomes (Van Rossum et al., 2000; Pfister and Gerstner, 2006). While tonic and burst firing are often studied in isolation, biological networks frequently alternate between them or express both within the same population, as seen in thalamic relay cells, hippocampal pyramidal neurons, and central pattern generators (McGinley et al., 2015; Sherman, 2001). The interaction between these modes in shaping long-term synaptic organization remains largely unexplored, both experimentally and computationally. Our model provides a controlled framework to map how each regime influences the geometry of weight space.

Our results show that collective burst firing drives synaptic weights toward a narrow region in weight space—a burst-induced attractor—whereas tonic firing maintains a more dispersed distribution. This convergence arises from synchrony-induced similarity in calcium dynamics or spike-time correlations across synapses, constraining their trajectories toward a shared attractor (Fig 5A). In contrast, tonic firing generates more heterogeneous activity patterns and multiple, separated convergence points. The mechanism is general, emerging across calcium-based and spike-based plasticity rules (Graupner and Brunel, 2012; Graupner et al., 2016; Pfister and Gerstner, 2006; Song et al., 2000; Deperrois and Graupner, 2020).

**Fig. 5:**
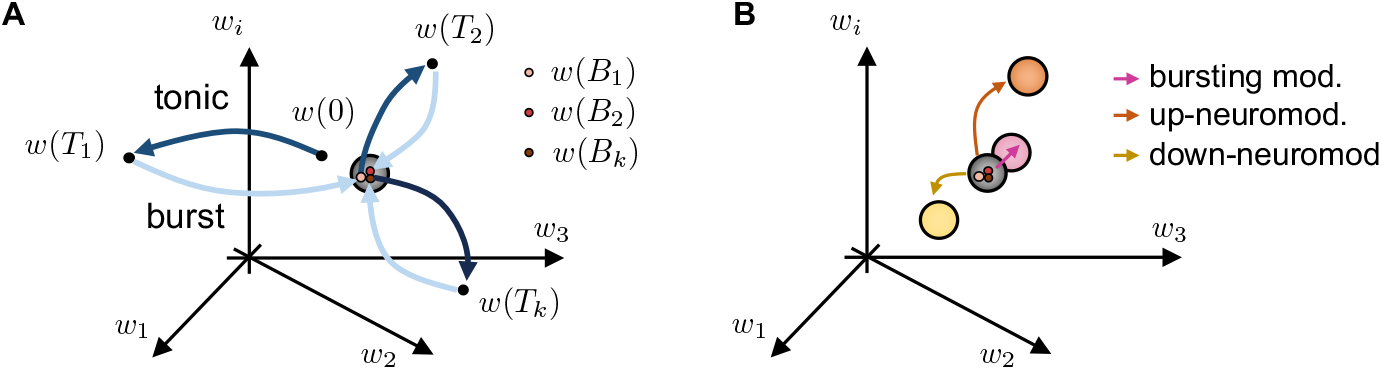
Conceptual illustration of the burst-induced attractor and its modulation. **A**. Example trajectories in synaptic weight space during transitions between tonic (light blue) and burst firing (dark blue). Tonic firing leads to more dispersed convergence points *w*(*T*_*k*_), whereas burst firing drives weights toward a narrower attractor region *w*(*B*_*k*_). **B**. Neuromodulation shifts the location of the burst-induced attractor; by modulating the collective burst firing (pink, referring to Fig 3, or by modulating the plasticity parameters in a weight-dependent manner mimicking synaptic tag (referring to Fig 4). Up-neuromodulation moves the attractor toward higher weights (orange), while down-neuromodulation shifts it toward lower weights (yellow).

We further show that the location of the burst-induced attractor depends on both neural dynamics and the parameters controlling potentiation and depression. Neuromodulatory changes to these parameters can shift the attractor, enabling global reorganization during rhythmic bursting (Frémaux and Gerstner, 2015; Brzosko et al., 2017; Seol et al., 2007; Pawlak, 2010). Implementing a tag-dependent rule, in which strong and weak synapses follow distinct neuromodulated kernels, creates two coexisting attractors: one stabilizing strong synapses and another downscaling weaker ones. This mechanism parallels the synaptic tagging and capture framework (Frey and Morris, 1997; Redondo and Morris, 2011), offering a computational route for selective long-term consolidation (Fig 5B). More generally, because the attractor is determined by plasticity parameters, neuromodulation and tagging can generate not only two, but potentially multiple stable consolidation attractors, allowing different synaptic selection profile.

An interesting extension is to incorporate additional hidden synaptic variables, such as eligibility traces or biochemical tags, which can store information independently of the expressed weight (Fusi et al., 2005; Benna and Fusi, 2016; Zenke et al., 2015; Gerstner et al., 2018). In such models, burst firing could pull the primary weight into the attractor while using hidden traces to transfer the acquired learning, allowing the coexistence of flexibility and long-term stability.

Overall, we identify burst-induced attractors as an emergent property of collective bursting combined with soft-bound plasticity rules. By demonstrating how neuromodulation and tagging can shift or split these attractors, we outline a computational pathway for synaptic selection and consolidation. Extending this framework with hidden synaptic traces may help reconcile the trade-off between adapting to new inputs and preserving past memories—a balance that biological networks likely exploit when alternating between tonic and burst activity states. This work finally suggests that similar principles may extend to other firing regimes.

## 6 Methods

All original data in this work were generated programmatically using the Julia programming language (Bezanson et al., 2017). Analyses were performed in Matlab. The code files are freely available at https://github.com/KJacquerie/Burst-Attractor. Any additional information required to reanalyze the data reported in this paper is available from the lead contact upon request.

### 6.1 Method details

#### 6.1.1 Neuron and network model

All neurons are modeled using a single-compartment conductance-based using Hodgkin and Huxley formalism (Hodgkin and Huxley, 1952). The membrane voltage *V* of a neuron evolves according to the equation described by (Drion et al., 2018; Jacquerie and Drion, 2021):

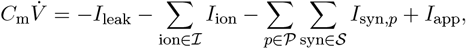

where *C*_m_ represents the membrane capacitance, *I*_leak_ is the leak current, *I*_ion_ are the intrinsic ionic currents with ℐ the set of all ionic channels, *I*_syn,*p*_ are the synaptic currents with the set of all synaptic neurotransmitter types and 𝒫 the set of all presynaptic neurons, and *I*_app_ denotes the applied current. Details about the ionic and synaptic currents and their associated dynamics and parameters are provided in Text S1.

The neuronal network comprises an inhibitory neuron ( 𝒩_inh_) projecting onto all excitatory neurons through GABA_A_ and GABA_B_ connections (refer to Fig 1B). Excitatory neurons are interconnected via a feedforward AMPA synapse, where the presynaptic neurons ( 𝒩_pre_) influence the postsynaptic neurons (𝒩 _post_). The number of excitatory neurons varies across different computational experiments. The excitatory synaptic current perceived by the postsynaptic neuron *i* from presynaptic neuron *j* is characterized by:

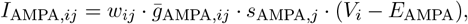

where *w*_*ij*_ represents the synaptic weight, 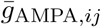 represent the maximal conductance of the AMPA postsynaptic receptor (AMPAr).The variable *s*_AMPA,*j*_ denotes the gating variable of the AMPAr, dynamically modulated by the presynaptic membrane voltage (*V*_*j*_) and *E*_AMPA_ is the reversal potential of AMPAr.

#### 6.1.2 Switch from tonic to burst firing

The inhibitory neuron dynamically influences the network activity. An external current applied to the inhibitory neuron (*I*_app,inh_) models the effects of a neuromodulatory signal. A depolarizing current induces tonic firing activity in this inhibitory neuron, causing all excitatory cells to remain at rest. A sufficiently large external depolarizing pulse can evoke action potentials in the excitatory neurons. Conversely, a hyperpolarizing current applied to the inhibitory neuron switches the entire network into a synchronized collective bursting activity throughout the network (Zagha and McCormick, 2014; Jacquerie and Drion, 2021). In each computational experiment, the depolarizing current *I*_app,inh_ is equal to 3 nA*/*cm^2^ for tonic state and −1.2 nA*/*cm^2^ for burst state.

#### 6.1.3 External applied pulse train current

In tonic firing states, each excitatory neuron *i* is triggered by an applied current (*I*_app,*i*_) to make the neuron fire at a nominal frequency *f*_0_. To generate this input-driven tonic activity, a neuron receives a pulse train current, where each pulse lasts 3 ms and has an amplitude that is independently sampled from a uniform distribution on an interval between 50 nA*/*cm^2^ and 60 nA*/*cm^2^. The interpulse intervals (*i*.*e*., the time between two successive pulses) are independently sampled from a Normal distribution with a mean equal to 1*/f*_0_ and a standard deviation equal to 0.1*/f*_0_.

In burst firing states, each excitatory neuron *i* undergoes a noisy baseline such as the applied current (*I*_app,*i*_) randomly fluctuating between the interval 0 and 1 nA*/*cm^2^.

#### 6.1.4 Homogeneous and heterogeneous network

We define a *homogeneous* network (or circuit), where each neuron has the same intrinsic ion channel maximal conductances 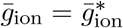 (numerical values provided in Text S1). We define a *heterogeneous* network (or circuit), where the maximal conductance 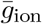 for each neuron is randomly chosen within a *±*10 % interval around its nominal value 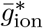, such as 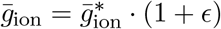 with *ϵ* ∼ Unif(−0.1, 0.1).

#### 6.1.5 Local field potential

The local field potential (LFP) measures the average behavior of interacting neurons. It reflects the collective excitatory synaptic activity received by the postsynaptic neuron population. The overall synaptic activity is measured by the mean of the individual synaptic currents:

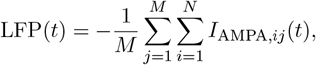

where *M* is the number of postsynaptic neurons and *N* is the number of presynaptic neurons (Jacquerie and Drion, 2021; Drion et al., 2018).

#### 6.1.6 Synaptic plasticity

##### Calcium-based model

The change in the synaptic weight *w*_*ij*_ between a presynaptic neuron *j* and a postsynaptic neuron *i* is governed by the calcium-based model proposed by (Graupner and Brunel, 2012):

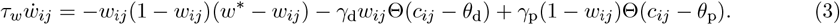

Here, *τ*_*w*_ represents the time constant, *w*^*^ defines the stable state (equal to 0.5), *γ*_p_ is the potentiation rate, *γ*_d_ is the depression rate, *θ*_p_ is the potentiation threshold, and *θ*_d_ is the depression threshold. The function Θ(· ) is the Heaviside function, which returns 1 if *c*_*ij*_ *> θ*_p_ and 0 otherwise.

The change in synaptic weight depends on the calcium concentration *c*_*ij*_, which is the sum of the calcium caused by the activity of the presynaptic neuron *j* and the activity of the postsynaptic neuron *i*. A pre-or postsynaptic spike translates into a calcium exponential decay. Further explanations are provided in Text S2. The calcium-based rule defined by (Graupner and Brunel, 2012) implements a soft-bound, where the perceived potentiation rate *γ*_p_(1 − *w*_*ij*_) is smaller for high *w*_*ij*_ than for low *w*_*ij*_. The same reasoning applies to the depression rate *γ*_d_. Detailed parameter values are provided in Text S3.

In Fig 2-4, the calcium-based model is modified according to the version of (Graupner et al., 2016). The model is similar except that the stable state is removed and the fitted plasticity parameters are adapted:

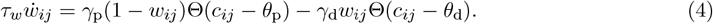

The only difference between the two calcium-based models is that when calcium levels fall below the depression threshold, the synaptic weight in the 2016 model is no longer subject to the cubic term and therefore remains unchanged. Both rules can be applied, and in either case the burst-induced attractor is present. However, the 2016 formulation is more convenient for demonstrating this phenomenon.

#### 6.1.7 Computational experiment related to Fig 1

Fig 1A-B illustrate the activity in the feedforward network, comprising 50 presynaptic neurons connected to 50 postsynaptic neurons. The network is heterogeneous. The network is in tonic firing mode during 1.5 s (see subsubsection 6.1.2). Each excitatory neuron receives a train of current pulses with nominal frequencies *f*_0_ randomly sampled from a uniform distribution between 0.1 and 50 Hz (see subsubsection 6.1.3). Then, the network switches to synchronized collective bursting during 2.5 s.

Fig 1C illustrates the evolution of the weights, consisting of 50 presynaptic neurons connected to 50 postsynaptic neurons, along with one inhibitory neuron projecting GABA current onto all of them.

It comprises a total of 2500 weights. This computational experiment consists of 8 states interleaving tonic and burst firing, each lasting 20 s. During tonic firing, neurons are stimulated with a pulse train current, where the pulse frequency for each neuron is randomly chosen from a uniform distribution between 0.1 Hz and 50 Hz.

Fig 1D demonstrates the values of the normalized weights at the end of the fourth tonic state (vertical axis) with the values of the normalized weights at the end of the third tonic state (horizontal axis). The weights are normalized using the formula

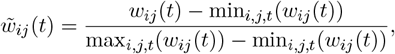

where max_*i,j,t*_(*w*_*ij*_(*t*)) and min_*i,j,t*_(*w*_*ij*_(*t*)) denote the maximum and minimum values obtained by comparing the values at the end of the third and fourth tonic states. The distribution of the normalized weights is represented for bins of width 0.025 between 0 and 1. Similarly, for burst firing (Fig 4E), the same data analysis protocol is used for the synaptic weights at the end of the third burst state and the end of the fourth burst state. The scatter plots computed for the other states are shown in Fig S1.

#### 6.1.8 Computational experiment related to Fig 2

Fig 2A-C illustrate the calcium fluctuations between two presynaptic neuron *j* connected to one postsynaptic neuron *i*. The calcium dynamics are following (Graupner and Brunel, 2012; Graupner et al., 2016) (see Text S2). Fig 2D-E are the time spent in each calcium region (see 4.5 for calculation). Fig 2F-H illustrate the evolution of the synaptic weights during two states of 50 s within the same network of 50 pre- and 50 postsynaptic neurons for 100 weights among 2500. The network has the same set of intrinsic and synaptic conductances as in Fig 1. The two states are initialized with the same set of initial conductances chosen randomly within 0 and 1. Fig 2G-I are the comparison of the predicted value given by the Equation 1 and the simulated value directly extracted during the simulation.

#### 6.1.9 Computational experiment related to Fig 3

Fig 3A (bottom) illustrates the evolution of the synaptic weights during four states of 20 s within the same network of 50 pre- and 50 postsynaptic neurons, along with one inhibitory neuron, maintaining identical maximal conductances as in Fig 1. The first state corresponds to a tonic state, where the inhibitory neuron is depolarized at 3 nA*/*cm^2^ during 20 s. The excitatory neurons receive a pulse train current with the pulse frequency randomly chosen in a uniform distribution between 0.1 Hz and 50 Hz for each neuron. This tonic state is succeeded by three bursting states, each lasting 20 s. The hyperpolarizing current successively equals -1.4, -1.6, and −1 nA*/*cm^2^. Fig 3A (top) presents the network activity and the associated LFP during 4s extracted from the middle of each state.

Fig 3B displays the distributions of the synaptic weights at the end of each state. The synaptic weights are not normalized, and the distribution is computed for bins of width 0.025 between 0 and 1.

#### 6.1.10 Computational experiment related to Fig 4

Fig 4B shows the evolution of synaptic weights over four 30 s states in six homogeneous circuits. Each circuit consists of one inhibitory neuron connected to two excitatory neurons. During the tonic firing states, three circuits are configured with correlated excitatory activity, where the pulse frequency is selected between 30 Hz and 60 Hz, and three circuits are configured with uncorrelated activity, with pulse frequencies between 0 Hz and 10 Hz. This setup allows us to sample a broad range of initial weight trajectories, from potentiation to depression. In the second burst state, the potentiation level Ω_p_ is reduced from 0.65 to 0.2.

Fig 4D uses the same circuit configuration and parameters, except that at the end of the second tonic state, synapses are tagged according to their weight values. Synapses with *w >* 0.75 at the end of the tonic state are assigned an up-neuromodulated calcium-based rule with Ω_p_ = 0.95, while the remaining synapses follow a down-neuromodulated rule with Ω_p_ = 0.2.

## Supplementary Material

### Text S1 Conductance-based model description

#### Neuron

The membrane voltage of the neuron is described by the Hodgkin and Huxley formalism such as:

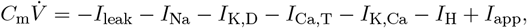

where

- 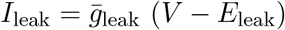 is a leaky current;
- 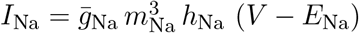 is a transient sodium current;
- 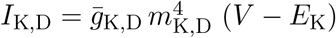 is a delayed-rectifier potassium current;
- 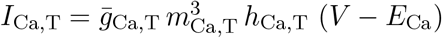 is a T-type calcium current;
- 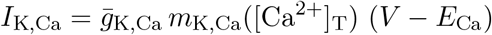 is a calcium-activated potassium current;
- 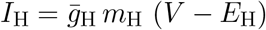 is a hyperpolarization-activated cation current;
- *I*_app_ is an applied current.

The membrane capacitance (expressed in µF*/*cm^2^) is *C*_m_ = 1, the reversal potentials (expressed in mV) are *E*_leak_ = −55, *E*_Na_ = 50, *E*_K_ = −85, *E*_Ca_ = 120, *E*_H_ = 20.

The ion channel maximal conductances (expressed in mS*/*cm^2^) are 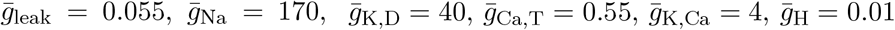.

The variables *m*_ion_ (resp. *h*_ion_) represents the activation (resp. inactivation) variable of the ion channel “ion”. Their dynamics are given by

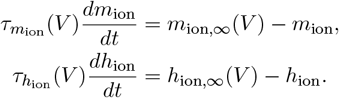

The steady-state values *x*_ion, ∞_ (*V* ) and the time constants 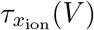 of the ion channel “ion” are voltage-dependent such as:

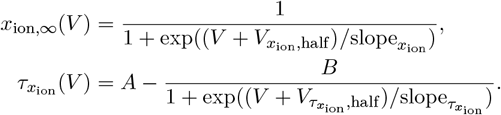

The parameters for the different channels are given following the format 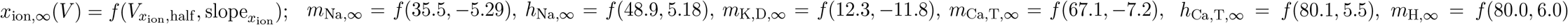 and 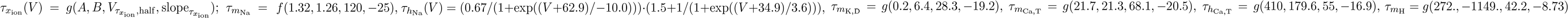.

In this conductance-based model, the calcium Ca^2+^ is entering through the T-type calcium channel.

The dynamics of the calcium concentration is thus given by:

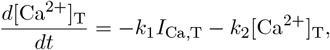

where *k*_1_ and *k*_2_ are the rate variables. The kinetics for the synapse are *k*_1_ = 0.1 M/msA, *k*_2_ = 0.01 nM/ms. The calcium-activated potassium current considers this calcium entry to update its gating variable:

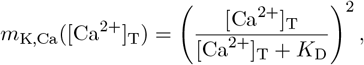

where *K*_D_ is a calcium-activation constant. The calcium half-activation constant is *K*_D_ = 170nM.

#### Network

We have three sets of neurons 𝒩_inh_, 𝒩_pre_, and 𝒩_post_. The excitatory synaptic current perceived by the postsynaptic neuron *i* from presynaptic neuron *j* is characterized in the main methods. The inhibitory synaptic current perceived by the postsynaptic neuron *i* from presynaptic neuron *j* is characterized by

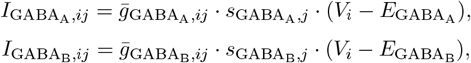

where 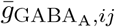 and 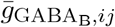 respectively represent the maximal conductance of the GABA_A_ receptors and the GABA_B_ receptors. Without variability, they are set to 2 mS*/*cm^2^ and 1.5 mS*/*cm^2^ . The variable 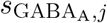 denotes the gating variable of the GABA_A_ postsynaptic receptor (GABA_A_r), dynamically modulated by the presynaptic membrane voltage (*V*_*j*_) and 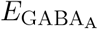 is the reversal potential of GABA_A_r (set to *E*_Cl_ = −70 mV)). The variable 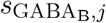 is the same for the the GABA_B_ postsynaptic receptor (GABA_B_r) where the reversal potential of GABA_B_r is set to *E*_K_ = −85 mV.

The gating variables of the synapses are 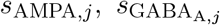 and 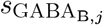 are variables whose dynamics depends on the considered presynaptic membrane potential following the equations

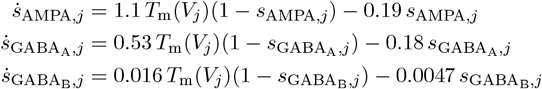

with transmitter concentration follows the formalism described in (Destexhe et al., 1994), that is,

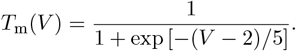

The numerical values (1.1, 0.19, 0.53, 0.18, 0.016, and 0.0047) are rate parameters to mimic the kinetics of the synaptic receptors. These values originated from (Destexhe et al., 1994). The smallest, the slowest the kinetics.

### Text S2 Plasticity implementation

In this section, the different plasticity rules and their variations are defined. The parameters are given in the code available on the GitHub https://github.com/KJacquerie/Burst-Attractor.

#### Calcium-based models

##### Calcium dynamics

We consider the linear calcium dynamics suggested by (Graupner and Brunel, 2012; Graupner et al., 2016). The presynaptic and postsynaptic spike contributions add linearly. Indeed, calcium enters from NMDA receptors and voltage-dependent calcium channels. Instead of describing the whole calcium, the phenomenological effect on the calcium variation is considered. At each pre or postsynaptic spike (respectively named as *t*_*j,k*_ and *t*_*i,k*_), the calcium immediately rises and then exponentially decays characterized by a calcium decay time constant equal to *τ*_Ca_:

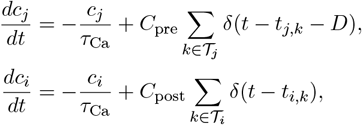

where *C*_pre_ and *C*_post_ are the presynaptically and postsynaptically evoked calcium amplitudes. The parameter *D* is a time-delay between the presynaptic spike and the corresponding postsynaptic calcium transient occurrence accounts for the slow rise time of the NMDAr-mediated calcium influx (Graupner and Brunel, 2012; Graupner et al., 2016).

The total calcium amplitude *c*_*ij*_(*t*) driving the synaptic change is given by:

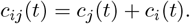

The time-evolution for several pre- and postsynaptic spiking activity is written such as (Graupner et al., 2016):

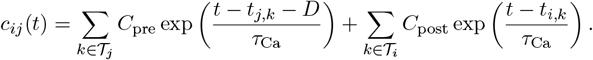

The resting calcium concentration is set to zero. The calcium concentrations are dimensionless. Both simplification is acknowledged because the synaptic rules are adapted in accordance. If a resting calcium concentration is wanted, the thresholds of potentiation and depression will be adapted. This notation follows the original paper notation.

##### [Graupner et al. 2012]

Graupner and Brunel developed a breakthrough calcium-dependent synaptic rule. It implements two opposing calcium-triggered pathways leading to an increase or a decrease in synaptic strength. Indeed, potentiation and depression are activated above calcium thresholds. It is defined by Equation 3

The parameters are fitted for the frequency plasticity-induced protocol (Sjöström et al., 2001) in the cortex (CTX) or the spike-time dependent plasticity-induced protocol (Bi and Poo, 1998) in the hippocampus (HPC).

##### [Graupner et al. 2016]

This model evolves in (Graupner et al., 2016) such as the synaptic plasticity rule becomes Equation 4

The parameters are fitted on CTX data in (Graupner et al., 2016). We have also matched the plasticity parameters to reproduce the data obtained in HPC.

This synaptic rule can be easily written using a standard form for ordinary differential equations with a steady-state value and a time constant for each calcium region as shown in Equation 5

##### [Shouval et al. 2012]

In (Shouval et al., 2002), the synaptic change follows a typical first-order differential equation:

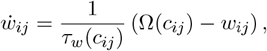

where the time constant *τ*_*w*_(*c*_*ij*_) and the steady-state value Ω(*c*_*ij*_) are calcium-dependent. This steady state value is defined by two sigmoids in order to build the U-shape for Ω:

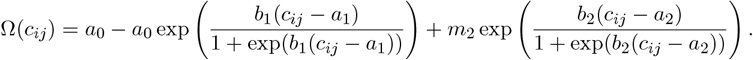

This expression relies on five parameters:

- *a*_0_ is the ordinate at low levels of calcium;
- *a*_1_ is the abscissa where the ordinate *a*_0_ is divided by 2 (it dictates the place along the x-axis where the first sigmoid is decreasing);
- *b*_1_ governs the sharpness of the decrease around *a*_1_ (the bigger, the flatter the slope);
- *m*_2_ the converging value at high calcium level;
- *a*_2_ is the x-value where Ω equal *m*_2_*/*2;
- *b*_2_ is similar as *b*_1_ (it dictates the sharpness of the slope).

The time constant is also calcium-dependent and it is given by (Shouval et al., 2002)

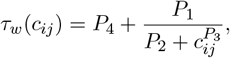

where *P*_1_, *P*_2_, *P*_3_, and *P*_4_ are fitted parameters to describe the calcium-dependent time constant. The different parameters were identified to reproduce the different potentiation and depression levels provided by the previous model for the frequency-dependency protocol (CTX, soft bounds). The synaptic rule is using the calcium dynamics presented above. The parameters are *a*_0_ = 0.5, *a*_1_ = 1.31, *a*_2_ = 1.8, *b*_1_ = 20, *b*_2_ = 40, *P*_1_ = 4*e*3, *P*_2_ = *P* 1.1*e* − 6, *P*_3_ = 2.4, *P*_4_ = 1. For hard bounds, the parameters become *m*_1_ = 0.25, *a*_1_ = 1, *a*_2_ = 2, *b*_1_ = 40, *b*_2_ = 10, *m*_2_, *P*_1_, *P*_2_, *P*_3_, *P*_4_ do not change.

##### [Deperrois et al. 2020]

The synaptic weight influences the postsynaptic calcium dynamics as mentioned in (Deperrois and Graupner, 2020). It is translated by a linear scale coupling between the presynaptically induced calcium amplitude:

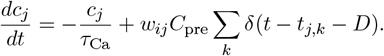

The effect of the short-term depression (STD) at the presynaptic site as described in (Deperrois and Graupner, 2020) is also tested in our project. To account for the available presynaptic resources, (Deperrois and Graupner, 2020) uses a variable *x*. Then, *U* considers the fraction of the resources required at each presynaptic spike and *τ*_rec_ is the resource recovery time constant for the variable *x* to come back to its resting state equal to 1

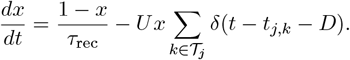

So we have

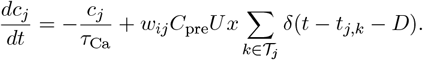

The parameters are fitted for the frequency-dependent plasticity-induced protocol (CTX) in soft or hard bounds, without short-term depression *τ*_*Ca*_ = 32.19, *C*_pre_ = 1.61, *C*_post_ = 1.124, *D* = 5.7527, *τ*_*w*_ = 79 975 ms, *θ*_p_ = 1.63, *θ*_d_ = 1, *γ*_p_ = 161.99, *γ*_d_ = 31.976; with short-term depression *τ*_*Ca*_ = 38.35, *C*_pre_ = 3.99, *C*_post_ = 1.29, *D* = 9.24, *τ*_*w*_ = 299 877.8 ms, *θ*_p_ = 1.63, *θ*_d_ = 1, *γ*_p_ = 564.4, *γ*_d_ = 111.3, *τ*_rec_ = 148.92, and *U* = 0.3838.

##### Hard-bound implementation

As pioneered in (Shouval et al., 2002), calcium drives the synaptic change. The first implementation suggested was:

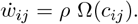

The synaptic change follows the speed given by Ω. It leads to weight runaway. The simplest solution to overcome this runaway is the addition of “hard bound” to constrain the synaptic weight between lower and an upper limit. In this work, they are respectively fixed at 0 and 1. Computationally, it is implemented such as: if(*w*_*ij*_ ≥ 1) : *w*_*ij*_ = 1; if(*w*_*ij*_ ≤ 0) : *w*_*ij*_ = 0.

We transformed the two-thresholds model suggested by (Graupner et al., 2016). The potentiation and depression terms (*γ*_p_ and *γ*_d_) are dependent on the synaptic weights. In phenomenological models, it is called ‘soft bounds’. A strong weight has a weaker effective potentiation rate and a stronger effective depression rate. By contrast, a weak weight has a stronger effective potentiation rate and a weaker depression rate. Mathematically, it is easily observed from the equation: 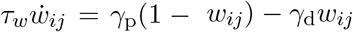. For a strong weight equal to 0.9: the expression becomes: 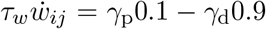. The effective potentiation rate is 10% the original value while the effective depression rate is 90 % is the original value.

To overcome this weight dependency, we convert the soft bounds expression into hard bounds:

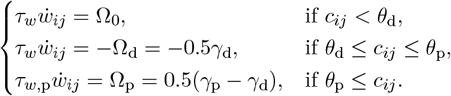

At low levels of calcium, the synaptic weight is unchanged (Ω_0_ = 0). At intermediate levels, we used the expression provided in the original model by considering a fixed mean weight equal to 0.5. Therefore, the synaptic change is decreased by a depression rate of 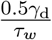. At high levels of calcium, the potentiation rate equals 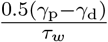. This transformation is valid because the provided potentiation rate is bigger than the depression rate (parameters originate from the paper). The presence of 0.5 comes from the removal of *w*-dependency in the main equation. To do so, its medium value has been validated during the fitting protocol in experimental data (Sjöström et al., 2001). Keeping *γ*_d_ for the depression rate and *γ*_p_ for the potentiation rate leads to a discrepancy with the experimental data mentioned.

The same development is performed for (Shouval et al., 2002). The synaptic rule simply becomes: 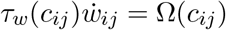 with the addition of hard bounds. Similar mathematical manipulations have been done in calcium-based rules established by considering the presynaptic resource depletion. The weight dependency has been removed inside the synaptic rule and the parameters have been updated. The parameters were once again fitted to reproduce the frequency-dependency protocol from (Sjöström et al., 2001).

#### Spike-time dependent models

##### Pair-based model

To reproduce the classical STDP window, pair-based model is implemented using synaptic traces, respectively *x* and *y* for the presynaptic neuron *j* and postsynaptic neuron *i*:

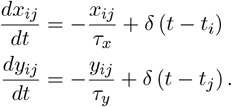

The weight change relative to the STDP window was then computed using the following equations, with explicit hard bounds defined such as 0 *< w*_*ij*_ *<* 1:

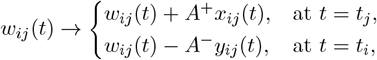

or using soft bounds:

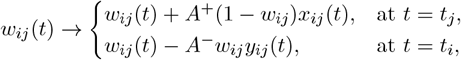

where *A*^+^ *>* 0, *A*^−^ *>* 0 and *t*_*i*_ (resp. *t*_*j*_) representing the spike timing of the presynaptic (resp. postsynaptic) neuron, *i*.*e*., each time *t* = *t*_*i*_ (resp. *t* = *t*_*j*_), we consider a post-pre (resp. pre-post) pair for the weight change (Morrison et al., 2008; Song et al., 2000; Van Rossum et al., 2000). In this project, we combine soft-bound and hard-bound dependency using the notation proposed in (Gütig et al., 2003):

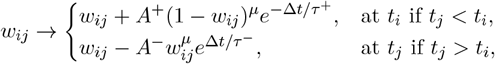

where *µ* is equal to 0 for hard-bounds and 1 for soft-bounds.

The parameters are *A*^+^=0.0096, *A*^−^=0.0053, *τ*_*x*_= 16.8 and *τ*_*y*_=33.7 (HPC) (Bi and Poo, 2001).

##### Triplet model

Similarly to the pair-based model, the triplet model was implemented using trace variables following (Pfister and Gerstner, 2006):

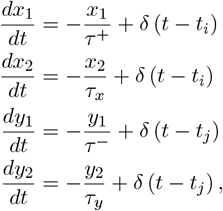

where *t*_*i*_ (resp. *t*_*j*_) is the timing of a presynaptic spike (resp. postsynaptic). The full model implemented by (Pfister and Gerstner, 2006) takes into account weight change due to pre-post, with the constant 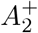, inducing potentiation or post-pre pairs, with the constant 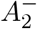, inducing depression (similar to classical pair-based model, with *x*_1_(*t*) and *r*_2_(*t*) as the presynaptic and postsynaptic traces, respectively with their respective time constant *τ* ^+^ and *τ*^−^).

The improvement over the classic pair-based model is that a triplet of spikes is also considered. Thanks to previously introduced traces *y*_2_, decaying with a time constant *τ*_*y*_, and *x*_2_, decaying with *τ*_*x*_, pre-post-pre triplets are treated (associated with the constant 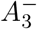, inducing depression) as well as post-pre-post triplets (associated with the constant 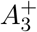, inducing potentiation).

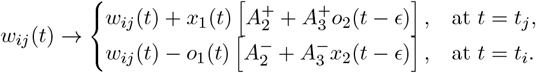

The parameters are for the minimal model (CTX) 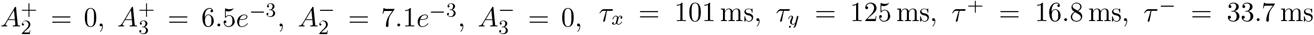 and (HPC) 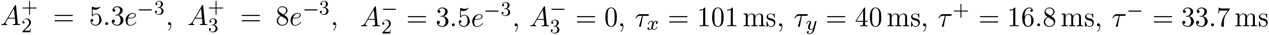.

The model can be described using soft bounds (Graupner et al., 2016):

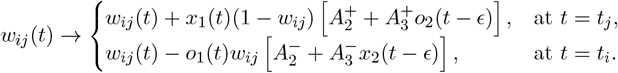

The parameters are (CTX) 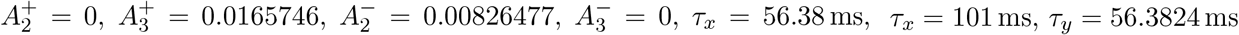, same *τ*^+^, *τ*^−^ and for (HPC) same as for hard-bounds.

### Text S3 Computational experiments: numerical values

The parameters associated with current and initial synaptic weights are constant in the different simulations: *I*_app,inh_(Tonic) = 3 nA*/*cm^2^, *I*_app,inh_ (Burst) = −1.2 nA*/*cm^2^, *w*_0_=0.5, 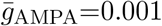.

The parameters associated with the synaptic plasticity are given in (Graupner and Brunel, 2012) fitting cortical (CTX) data (Sjöström et al., 2001): *τ*_Ca_ = 22.6936 ms, *C*_pre_ = 0.56, *C*_post_ = 1.24, *D* = 4.60 ms, *τ*_*w*_ = 346.3615 *×* 10^3^ ms, *γ*_p_ = 725.085 *×* 1.1 (Tonic), *γ*_p_ = 725.085 *×* 0.95 (Burst), *γ*_d_ = 331.909, *θ*_p_ = 1.3, *θ*_d_ = 1, *w*^*^ = 0.5. The potentiation rate *γ*_*p*_ is slightly scaled up during tonic firing to induce stronger potentiation compared to the initial model, and it is reduced by 5 % during burst firing to place the fixed-point at a lower value compared to the initial model.

For (Graupner and Brunel, 2012) fitting hippocampus (HPC) data (Bi and Poo, 1998): *τ*_Ca_ = 20 ms, *C*_pre_ = 1, *C*_post_ = 2, *D* = 13.7 ms, *τ*_*w*_ = 150 *×* 10^3^ ms, *γ*_p_ = 321.808 *γ*_d_ = 200, *θ*_p_ = 1.3, *θ*_d_ = 1, *w*^*^ = 0.5.

The parameters associated with the synaptic plasticity are given in (Graupner et al., 2016) fitting cortical (CTX) data (Sjöström et al., 2001): *τ*_Ca_ = 22.272 12 ms, *C*_pre_ = 0.8441, *C*_post_ = 1.62138, *D* = 9.537 09 ms, *τ*_*w*_ = 520 761.29 ms, *γ*_p_ = 597.08922, *γ*_d_ = 137.7586, *θ*_p_ = 2.009289, *θ*_d_ = 1. For (Graupner et al., 2016) in soft bounds that fit the hippocampal data of (Bi and Poo, 1998), the parameters remain the same, except that *θ*_*p*_ becomes equal to 1.45.

The parameters used in each simulation are for Fig 1 *N* = 50, *M* = 50, *T*_state_ = 20 s, *N*_state_ = 8, for Fig 2 *N* = 50, *M* = 50, *T*_state_ = 50 s, for Fig 3 *N* = 50, *M* = 50, *T*_state_ = 20 s, and for Fig 4 *N*_state_ = 4, *N* = 1, *M* = 1,*T*_state_ = 20 s .

### Text S4 Derivation of the burst-induced attractor in a calcium-based model

We consider the calcium-based plasticity model of Graupner and Brunel (Graupner et al., 2016), given in Equation 4. This rule specifies how the synaptic weight *w*_*ij*_ evolves as a function of the postsynaptic calcium concentration *c*_*ij*_. Depending on the calcium level, the dynamics are governed by three regimes: when calcium remains below the depression threshold (*c*_*ij*_ *< θ*_d_), the weight does not change; at intermediate calcium levels (*θ*_d_ ≤ *c*_*ij*_ *< θ*_p_), the weight relaxes toward the depression steady state Ω_d_ = 0 with time constant *τ*_*w*,d_ = *τ*_*w*_*/γ*_d_; and at high calcium concentrations (*c*_*ij*_ ≥ *θ*_p_), the weight relaxes toward the potentiation steady state Ω_p_ = *γ*_p_*/*(*γ*_p_ +*γ*_d_) with time constant *τ*_*w*,p_ = *τ*_*w*_*/*(*γ*_p_ +*γ*_d_). These regimes are summarized in:

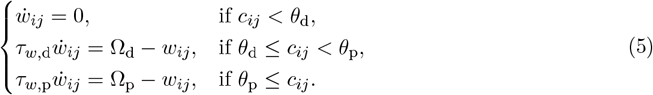

Because calcium fluctuates on the timescale of individual spikes and bursts (milliseconds), while synaptic weights evolve on much slower timescales (seconds to minutes), we reformulate the dynamics by averaging over an intermediate window of length *T* . The interval *T* must be long enough to capture typical calcium fluctuations. The averaged dynamics are then given by

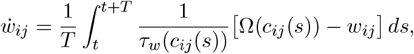

where Ω(*c*_*ij*_) denotes the steady-state value (either Ω_d_ or Ω_p_) associated with a given calcium regime, and *τ*_*w*_(*c*_*ij*_) is the corresponding time constant. This formulation describes the effective pull exerted on the synaptic weight by the sequence of calcium transients encountered during the interval.

To express this average more explicitly, we define the effective times spent in each regime. The effective depression time *α*_d,*ij*_ is the fraction of the interval spent between the depression and potentiation thresholds, normalized by *τ*_d_. Similarly, the effective potentiation time *α*_p,*ij*_ measures the fraction of time spent above the potentiation threshold, normalized by *τ*_p_:

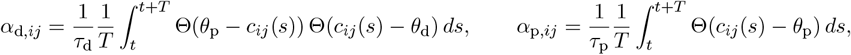

where Θ is the Heaviside step function. Periods where calcium remains below *θ*_d_ not contribute to a change and are quantified by *α*_0_. Fig 2A illustrates how these contributions are measured for a representative calcium trace.

With these definitions, the averaged dynamics is simplified to

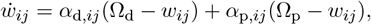

which makes explicit that the weight is simultaneously pulled toward the depression and potentiation steady states, with strengths proportional to the effective times spent in each regime. The fixed point 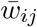 is reached when these contributions balance, yielding Equation 1. This expression shows that the long-term value of the synaptic weight is a weighted average of the potentiation and depression steady states, where the weights reflect how often calcium crosses each threshold (Gütig et al., 2003; Dorman and Blackwell, 2021). In other words, the precise statistics of calcium fluctuations determine the effective balance of potentiation and depression, and thus the fixed point to which the synapse converges.

### Text S5 Derivation of the burst-induced attractor in a spike-time dependent plasticity rule

We next derive the fixed point for a spike-timing–dependent plasticity (STDP) rule. The pair-based model considers presynaptic and postsynaptic spike times (resp. *t*_*i*_ and *t*_*j*_), with their relative timing with a time difference Δ*t* = *t*_*j*_ − *t*_*i*_ to induce the change in synaptic weight. A classical pair-based STDP window is used: when a presynaptic spike precedes a postsynaptic spike, the weight increases (potentiation), and when the order is reversed, the weight decreases (depression) (Abbott and Nelson, 2000; Song et al., 2000; Morrison et al., 2008; Rubin et al., 2001; Van Rossum et al., 2000):

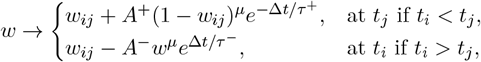

where *A*^+^ and *A*^−^ are the potentiation and depression parameters, *µ* stands for the weight-dependency, 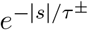 stands for the STDP kernel in potentiation or depression with *τ* ^+^ and *τ*^−^ being the time constants given in the pair-based model. The plasticity parameters *A*^+^ *>* 0, *A*^−^ *>* 0 (Morrison et al., 2008; Song et al., 2000). The weight dynamics can be constrained in two manners; either by using *hard bounds* or *soft bounds*. Hard bounds permit to stop the weight increase or decrease by adding upper or lower limits. Soft bounds decelerate the evolution if the weight reaches a bound. It is modeled by the weight-dependency parameter *µ*: *µ* is equal to 0 for hard-bounds and to 1 for soft-bounds) (Gerstner and Kistler, 2002).

The functions 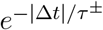 are the temporal kernel of potentiation and depression. If we introduce *S*_*i*_(*t*) = ∑_*k*_ *δ*(*t t*_*i,k*_) and *S*_*j*_(*t*) = ∑_*k*_ *δ*(*t t*_*j,k*_) for the spike trains of presynaptic neuron *j* and the postsynaptic neuron *i*, the evolution of the synaptic weight can be written as follows:

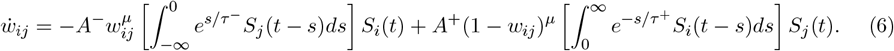

The time evolution of a weight and its convergence occurs on a time interval much larger than typical interspike intervals. Therefore, following the work of (Gütig et al., 2003; Legenstein and Maass, 2005), we can average the dynamics of the synaptic weight over a time interval *T* and get

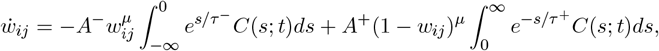

where *C*(*s*; *t*) is the (temporally averaged) correlation function between the pre and post spike trains, respectively noted *S*_*i*_(*t*) = ∑_*k*_ *δ*(*t* − *t*_*i,k*_) and *S*_*j*_(*t*) = ∑_*k*_ *δ*(*t* − *t*_*j,k*_), that is,

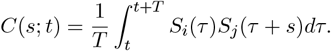

Assuming the stationary property of both spike trains is valid because the two neurons are bursting in a collective and synchronized manner. The correlation function becomes time-invariant, *i*.*e*., *C*(*s*; *t*) = *C*(*s*), and the time evolution of the synaptic weight can be computed as (Legenstein and Maass, 2005):

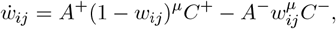

where 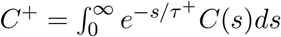 and 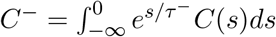.

A qualitative analysis of this equation helps to understand why synaptic weights converge towards a single steady-state for any stationary value of *C*(*s*), considering soft-bound dependency, *i*.*e*., *µ* = 1. The term *A*^+^(1 − *w*_*ij*_)*C*^+^ computes the weight increase due to all postsynaptic spikes following presynaptic spikes considering the associated time lag. The term *A*^−^*w*_*ij*_*C*^−^ computes the weight decrease due to all postsynaptic spikes preceding presynaptic spikes. When modeling soft bounds, both terms are weight-dependent, which deforms the plasticity kernel. When the synaptic weight is weak, the term *A*^+^(1 − *w*_*ij*_)*C*^+^ dominates, and potentiation overcomes depression. When the synaptic weight is strong, the term *A*^−^*w*_*ij*_*C*^−^ dominates, and depression overcomes potentiation. The drift eventually stabilizes at the synaptic weight value for which depression balances potentiation, *i*.*e*., *A*^+^(1 − *w*_*ij*_)*C*^+^ = *A*^−^*w*_*ij*_*C*^−^. This convergence value can be computed analytically by solving this balance equation.

The fixed-point value 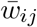 is obtained analytically by 2. This equation states that the fixed-point value depends on a ratio between the potentiation parameter weighted by the correlation between pre-post spikes and the depression parameter weighted by the correlation between post-pre spikes.

### Text S6 Derivation of the burst-induced attractor in a model using hard-bounds

The analytical analyses can be extended to the case of hard bounds. In this case, we consider the speed *λ*_*ij*_ of change, or slope. For the calcium-based model (Graupner et al., 2016), we obtain that slope corresponds to the sum of the depression rate and potentiation rate, each weighted by the fraction of time spent in their corresponding regions *α*_d_ and *α*_p_, which writes

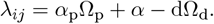

For the spike-time dependent plasticity model (Song et al., 2000), the slope can be predicted by the equation

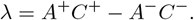

## Supplementary Figures

**Fig. S1:**
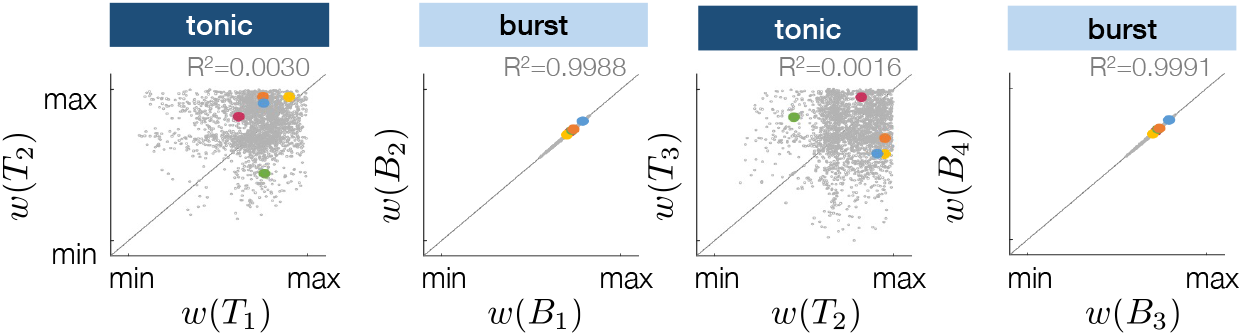
Replication of the analysis illustrated in Fig 1D-E at different tonic and burst states.

**Fig. S2:**
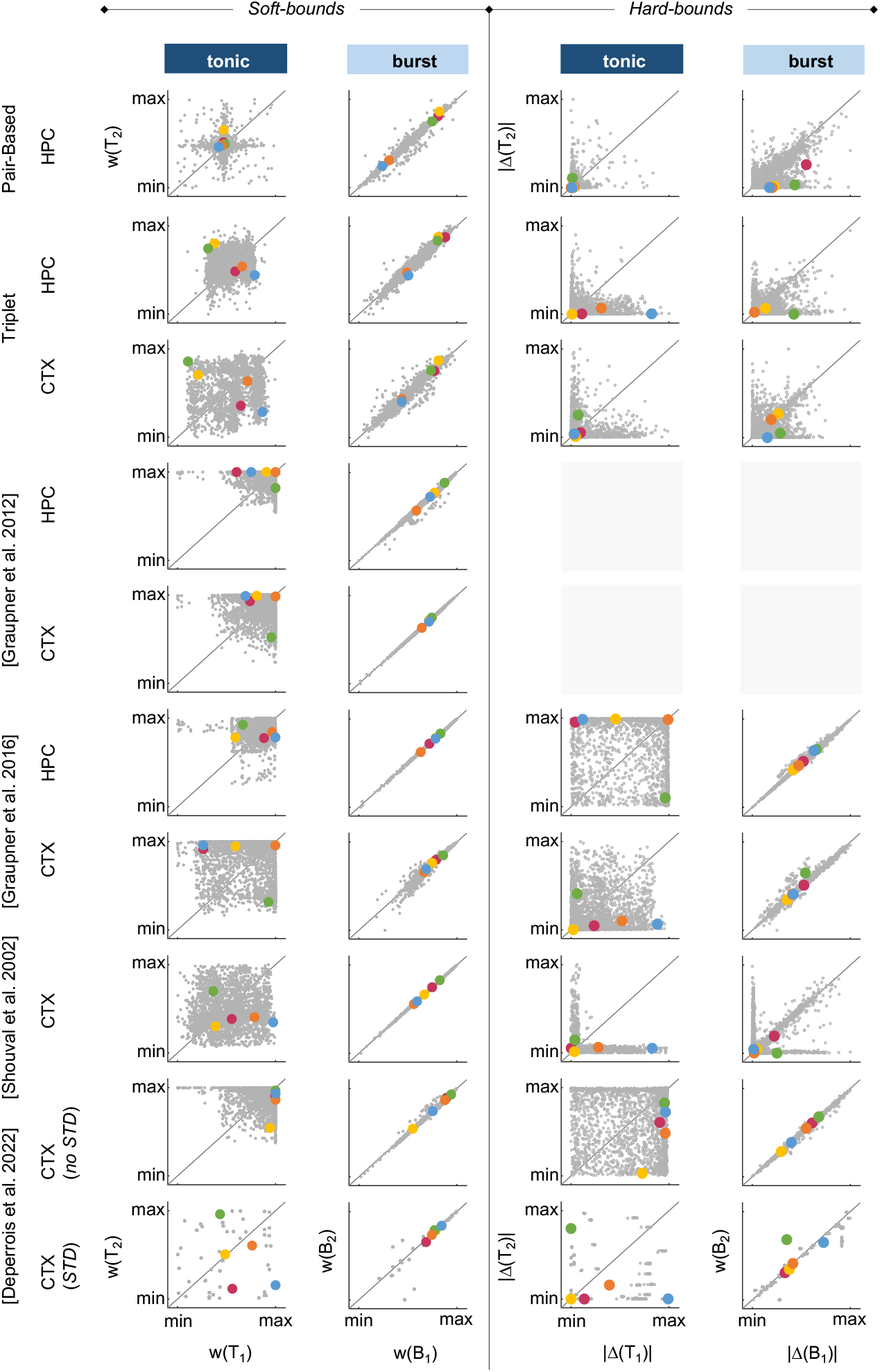
Replication of the experiment illustrated in Fig 1C in various synaptic plasticity rules using soft-bounds (left panel) and hard-bounds (right panel). Comparison of the synaptic weights at the end of the third and fourth tonic firing states (left column) or the third and fourth burst firing states (right column), normalized between the minimal and maximal values. (CTX=cortex, data fitted on (Sjöström et al., 2001); HPC=hippocampus, data fitted on (Bi and Poo, 1998)) Fig S3 shows the evolution of synaptic weights between two excitatory neurons during burst firing for different initial conditions (0:0.1:1). In blue, trajectories correspond to unmodulated plasticity parameters, while in yellow they show neuromodulated parameters. The color gradient emphasizes the initial strength of the synaptic weights, with darker shades indicating larger initial values.

**Fig. S3:**
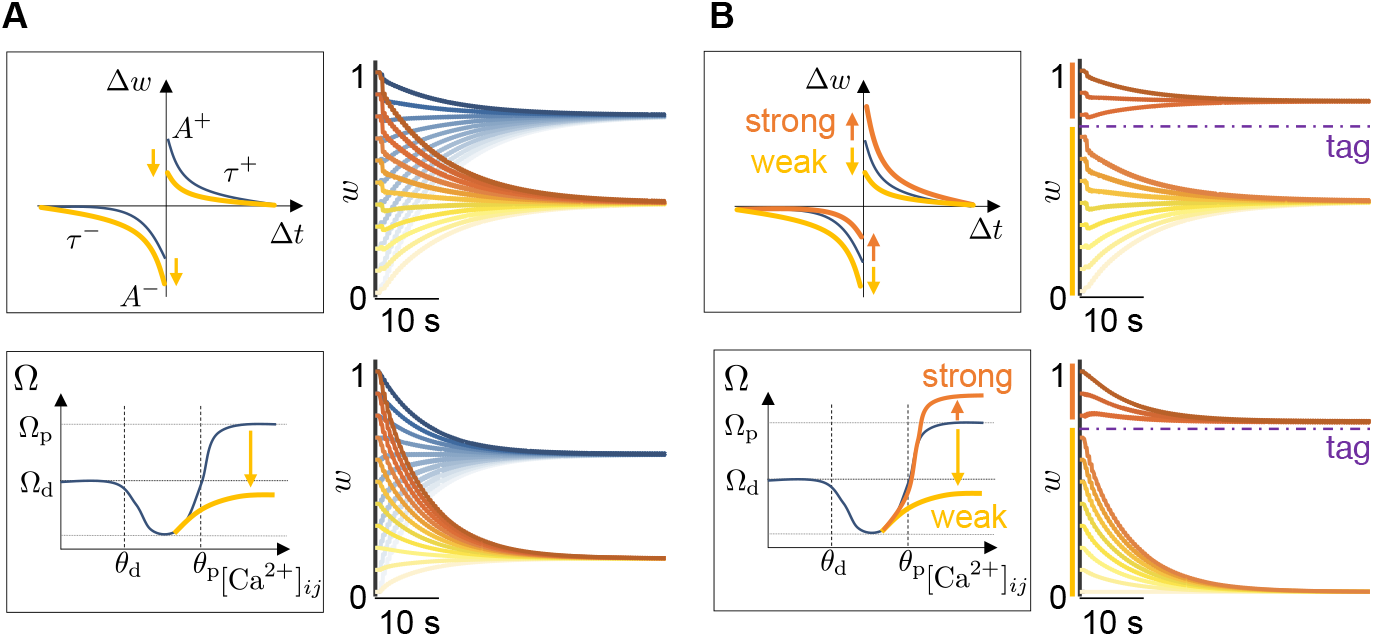
A. Global neuromodulation. Left: plasticity rules in spike-based (top) and calcium-based (bottom) formulations, where potentiation and depression parameters (*A*^*p*^,*A*^*m*^, or Ω_p_, Ω_d_) are globally downscaled (yellow arrows). Right: synaptic weights *w* converge toward a lower attractor during bursting, showing that global parameter changes shift the attractor in weight space. B. Tag-dependent modulation. Left: plasticity rules where synapses above a tagging threshold receive enhanced potentiation parameters, while weaker synapses are downscaled. Right: tagged synapses converge to a higher attractor (purple dashed line), while untagged synapses decay toward a lower one, resulting in bimodal consolidation. This mechanism shows how neuromodulation and tagging can selectively stabilize strong synapses while weakening weaker ones.

## Acknowledgments

Kathleen Jacquerie was a Research Fellow of the Fonds de la Recherche Scientifique—FNRS. This work was supported by the Belgian Government through the Federal Public Service Policy and Support. The authors thank Nora Benghalem, Juliette Ponnet, and Caroline Minne for their help in the early stages of the project. The authors acknowledge the valuable insight and feedback from Professor Eve Marder on the project.

## References

Abbott LF, Nelson SB (2000) Synaptic plasticity: Taming the beast. Nature Neuroscience 3:1178–1183.

Bazhenov M, Timofeev I, Steriade M, Sejnowski TJ (2002) Model of thalamocortical slow-wave sleep oscillations and transitions to activated states. Journal of Neuroscience 22:8691–8704.

Bazzari A, Parri H (2019) Neuromodulators and Long-Term Synaptic Plasticity in Learning and Memory: A Steered-Glutamatergic Perspective. Brain Sciences 9:300.

Benna MK, Fusi S (2016) Computational principles of synaptic memory consolidation. Nature Neuroscience 19:1697–1706.

Bezanson J, Edelman A, Karpinski S, Shah VB (2017) Julia: A Fresh Approach to Numerical Computing. SIAM Review 59:65–98.

Bi GQ, Poo MM (1998) Synaptic modifications in cultured hippocampal neurons: Dependence on spike timing, synaptic strength, and postsynaptic cell type. Journal of Neuroscience 18:10464–10472.

Bi Gq, Poo Mm (2001) Synaptic Modification by Correlated Activity: Hebb’s Postulate Revisited. Annual Review of Neuroscience 24:139–166.

Brzosko Z, Mierau SB, Paulsen O (2019) Neuromodulation of Spike-Timing-Dependent Plasticity: Past, Present, and Future. Neuron 103:563–581 Publisher: Elsevier Inc.

Brzosko Z, Zannone S, Schultz W, Clopath C, Paulsen O (2017) Sequential neuromodulation of Hebbian plasticity offers mechanism for effective reward-based navigation. eLife 6:e27756.

Butson CR (2012) Computational Models of Neuromodulation. International Review of Neurobiology 107:5–22.

Citri A, Malenka RC (2008) Synaptic plasticity: Multiple forms, functions, and mechanisms. Neuropsychopharmacology 33:18–41.

Dastgheib M, Kulanayagam A, Dringenberg HC (2022) Is the role of sleep in memory consolidation overrated? Neuroscience & Biobehavioral Reviews 140:104799.

Deperrois N, Graupner M (2020) Short-term depression and long-term plasticity together tune sensitive range of synaptic plasticity. PLoS Computational Biology 16:1–25.

Desroches M, Rinzel J, Rodrigues S (2022) Classification of bursting patterns: A tale of two ducks. PLOS Computational Biology 18:e1009752.

Destexhe A, Mainen ZF, Sejnowski TJ (1994) Synthesis of models for excitable membranes, synaptic transmission and neuromodulation using a common kinetic formalism. Journal of Computational Neuroscience 1:195–230.

Dorman DB, Blackwell KT (2021) Synaptic plasticity is predicted by spatiotemporal firing rate patterns and robust to in vivo-like variability preprint, Neuroscience.

Drion G, Dethier J, Franci A, Sepulchre R (2018) Switchable slow cellular conductances determine robustness and tunability of network states. PLoS Computational Biology 14:1–20.

Fauth M, Tetzlaff C (2016) Opposing Effects of Neuronal Activity on Structural Plasticity. Frontiers in Neuroanatomy 10.

Fernandez LM, Luthi A (2019) Sleep Spindles: Mechanisms and Functions. Physiological Reviews 100:805–868.

Foncelle A, Mendes A, Jedrzejewska-Szmek J, Valtcheva S, Berry H, Blackwell KT, Venance L (2018) Modulation of spike-timing dependent plasticity: Towards the inclusion of a third factor in computational models. Frontiers in Computational Neuroscience 12:1–21.

Frey U, Morris RGM (1997) Synaptic tagging and long-term potentiation. Nature 385:533–536.

Frémaux N, Gerstner W (2015) Neuromodulated spike-timing-dependent plasticity, and theory of three-factor learning rules. Frontiers in Neural Circuits 9.

Fusi S, Drew PJ, Abbott L (2005) Cascade Models of Synaptically Stored Memories. Neuron 45:599–611.

Gerstner W, Kistler WM (2002) Mathematical formulations of Hebbian learning. Biological Cybernetics 87:404–415.

Gerstner W, Lehmann M, Liakoni V, Corneil D, Brea J (2018) Eligibility Traces and Plasticity on Behavioral Time Scales: Experimental Support of NeoHebbian Three-Factor Learning Rules. Frontiers in Neural Circuits 12:1–16.

Gjorgjieva J, Toyoizumi T, Eglen SJ (2009) Burst-time-dependent plasticity robustly guides ON/OFF segregation in the lateral geniculate nucleus. PLoS Computational Biology 5.

González-Rueda A, Pedrosa V, Feord RC, Clopath C, Paulsen O (2018) Activity-Dependent Down-scaling of Subthreshold Synaptic Inputs during Slow-Wave-Sleep-like Activity In Vivo. Neuron 97:1244–1252.e5.

Graupner M, Brunel N (2012) Calcium-based plasticity model explains sensitivity of synaptic changes to spike pattern, rate, and dendritic location. Proceedings of the National Academy of Sciences of the United States of America 109:3991–3996.

Graupner M, Wallisch P, Ostojic S (2016) Natural firing patterns imply low sensitivity of synaptic plasticity to spike timing compared with firing rate. Journal of Neuroscience 36:11238–11258.

Gütig R, Aharonov R, Rotter S, Sompolinsky H (2003) Learning Input Correlations through Nonlinear Temporally Asymmetric Hebbian Plasticity. The Journal of Neuroscience 23:3697–3714.

Hill S, Tononi G (2005) Modeling sleep and wakefulness in the thalamocortical system. Journal of Neurophysiology 93:1671–1698.

Hodgkin AL, Huxley AF (1952) A quantitative description of membrane current and its application to conduction and excitation in nerve. The Journal of Physiology 117:500–544.

Izhikevich EM (2007) Solving the distal reward problem through linkage of STDP and dopamine signaling. Cerebral Cortex 17:2443–2452.

Jacquerie K, Drion G (2021) Robust switches in thalamic network activity require a timescale separation between sodium and T-type calcium channel activations. PLOS Computational Biology 17:e1008997.

Jedrzejewska-Szmek J, Damodaran S, Dorman DB, Blackwell KT (2017) Calcium dynamics predict direction of synaptic plasticity in striatal spiny projection neurons. European Journal of Neuroscience 45:1044–1056.

Kirkwood A (2007) Neuromodulation of Cortical Synaptic Plasticity In Tseng KY, Atzori M, editors, Monoaminergic Modulation of Cortical Excitability, pp. 209–216. Springer US, Boston, MA.

Lamprecht R, LeDoux J (2004) Structural plasticity and memory. Nature Reviews Neuroscience 5:45–54.

Lee SH, Dan Y (2012) Neuromodulation of Brain States. Neuron 76:209–222 Publisher: Elsevier Inc.

Legenstein RA, Maass W (2005) A Criterion for the Convergence of Learning with Spike Timing Dependent Plasticity In NIPS 2005, p. 8, Vancouver, Canada.

Lisman J, Grace AA, Duzel E (2011) A neoHebbian framework for episodic memory; role of dopamine-dependent late LTP. Trends in Neurosciences 34:536–547.

Magee JC, Grienberger C (2020) Synaptic Plasticity Forms and Functions. Annual Review of Neuroscience 43:95–117 Publisher: Annual Reviews.

McCormick DA, Salkoff DB (2015) Brain State Dependent Activity in the Cortex and Thalamus. Current Opinion in Neurobiology 31:133–140.

McGinley MJ, Vinck M, Reimer J, Batista-Brito R, Zagha E, Cadwell CR, Tolias AS, Cardin JA, McCormick DA (2015) Waking State: Rapid Variations Modulate Neural and Behavioral Responses. Neuron 87:1143–1161 Publisher: Elsevier.

Morrison A, Diesmann M, Gerstner W (2008) Phenomenological models of synaptic plasticity based on spike timing. Biological Cybernetics 98:459–478.

Nadim F, Bucher D (2014) Neuromodulation of neurons and synapses. Current Opinion in Neurobiology 29:48–56.

Okuda K, Højgaard K, Privitera L, Bayraktar G, Takeuchi T (2021) Initial memory consolidation and the synaptic tagging and capture hypothesis. European Journal of Neuroscience 54:6826–6849.

Olcese U, Esser SK, Tononi G (2010) Sleep and synaptic renormalization: A computational study. Journal of Neurophysiology 104:3476–3493.

Pawlak V (2010) Timing is not everything: neuromodulation opens the STDP gate. Frontiers in Synaptic Neuroscience 2.

Pedrosa V, Clopath C (2017) The role of neuromodulators in cortical plasticity. A computational perspective. Frontiers in Synaptic Neuroscience 8:38–38.

Pfister JP, Gerstner W (2006) Triplets of spikes in a model of spike timing-dependent plasticity. Journal of Neuroscience 26:9673–9682.

Redondo RL, Morris RGM (2011) Making memories last: the synaptic tagging and capture hypothesis. Nature Reviews Neuroscience 12:17–30.

Rubin J, Lee DD, Sompolinsky H (2001) Equilibrium Properties of Temporally Asymmetric Hebbian Plasticity. Physical Review Letters 86:364–367.

Salgado H, Köhr G, Treviño M (2012) Noradrenergic ‘Tone’ Determines Dichotomous Control of Cortical Spike-Timing-Dependent Plasticity. Scientific Reports 2:417.

Seibt J, Frank MG (2019) Primed to sleep: The dynamics of synaptic plasticity across brain states. Frontiers in Systems Neuroscience 13:1–19.

Seol GH, Ziburkus J, Huang S, Song L, Kim IT, Takamiya K, Huganir RL, Lee HK, Kirkwood A (2007) Neuromodulators Control the Polarity of Spike-Timing-Dependent Synaptic Plasticity. Neuron 55:919–929.

Sherman SM (2001) Tonic and burst firing: dual modes of thalamocortical relay. Trends in neurosciences 24:122–6.

Shine JM, Müller EJ, Munn B, Cabral J, Moran RJ, Breakspear M (2021) Computational models link cellular mechanisms of neuromodulation to large-scale neural dynamics. Nature Neuroscience 24:765–776.

Shouval HZ, Bear MF, Cooper LN (2002) A unified model of NMDA receptor-dependent bidirectional synaptic plasticity. Proceedings of the National Academy of Sciences of the United States of America 99:10831–10836.

Sjöström PJ, Turrigiano GG, Nelson SB (2001) Rate, Timing, and Cooperativity Jointly Determine Cortical Synaptic Plasticity. Neuron 32:1149–1164.

Song S, Miller KD, Abbott LF (2000) Competitive Hebbian learning through spike-timing-dependent synaptic plasticity. Nature Neuroscience 3:919–926.

Timofeev I, Grenier F, Bazhenov M, Houweling AR, Sejnowski TJ, Steriade M (2002) Short-and medium-term plasticity associated with augmenting responses in cortical slabs and spindles in intact cortex of cats in vivo. Journal of Physiology 542:583–598.

Tyree S, Luis De Lecea (2017) Optogenetic Investigation of Arousal Circuits. International Journal of Molecular Sciences 18:1773.

Tyulmankov D (2025) Computational models of learning and synaptic plasticity In Learning and Memory: A Comprehensive Reference, pp. 1–21. Elsevier.

Van Rossum MC, Bi GQ, Turrigiano GG (2000) Stable Hebbian learning from spike timing-dependent plasticity. Journal of Neuroscience 20:8812–8821.

Van Rossum MC, Shippi M (2013) Computational modelling of memory retention from synapse to behaviour. Journal of Statistical Mechanics: Theory and Experiment 2013.

Zagha E, McCormick DA (2014) Neural control of brain state. Current Opinion in Neurobiology 29:178–186 Publisher: Elsevier Ltd.

Zannone S, Brzosko Z, Paulsen O, Clopath C (2018) Acetylcholine-modulated plasticity in reward-driven navigation: a computational study. Scientific Reports 8:1–20.

Zeldenrust F, Wadman WJ, Englitz B (2018) Neural Coding With Bursts—Current State and Future Perspectives. Frontiers in Computational Neuroscience 12:48.

Zenke F, Agnes EJ, Gerstner W (2015) Diverse synaptic plasticity mechanisms orchestrated to form and retrieve memories in spiking neural networks. Nature Communications 6.

