## Supplementary Information for "Switching from tonic to burst firing creates an attractor in synaptic weight dynamics"

### Supplementary Material

#### Text S1 Conductance-based model description

**Neuron** The membrane voltage of the neuron is described by the Hodgkin and Huxley formalism such as:

$$C_\mathrm{m}\dot{V} = -I_\mathrm{leak} - I_\mathrm{Na} - I_\mathrm{K,D} - I_\mathrm{Ca,T} - I_\mathrm{K,Ca} - I_\mathrm{H} + I_\mathrm{app},$$

where

- $I_\mathrm{leak} = \bar{g}_\mathrm{leak}\,(V - E_\mathrm{leak})$ is a leaky current;
- $I_\mathrm{Na} = \bar{g}_\mathrm{Na}\,m_\mathrm{Na}^3\,h_\mathrm{Na}\,(V - E_\mathrm{Na})$ is a transient sodium current;
- $I_\mathrm{K,D} = \bar{g}_\mathrm{K,D}\,m_\mathrm{K,D}^4\,(V - E_\mathrm{K})$ is a delayed-rectifier potassium current;
- $I_\mathrm{Ca,T} = \bar{g}_\mathrm{Ca,T}\,m_\mathrm{Ca,T}^3\,h_\mathrm{Ca,T}\,(V - E_\mathrm{Ca})$ is a T-type calcium current;
- $I_\mathrm{K,Ca} = \bar{g}_\mathrm{K,Ca}\,m_\mathrm{K,Ca}([\mathrm{Ca}^{2+}]_\mathrm{T})\,(V - E_\mathrm{Ca})$ is a calcium-activated potassium current;
- $I_\mathrm{H} = \bar{g}_\mathrm{H}\,m_\mathrm{H}\,(V - E_\mathrm{H})$ is a hyperpolarization-activated cation current;
- $I_\mathrm{app}$ is an applied current.

The membrane capacitance (expressed in µF/cm$^2$) is $C_\mathrm{m} = 1$, the reversal potentials (expressed in mV) are $E_\mathrm{leak} = -55$, $E_\mathrm{Na} = 50$, $E_\mathrm{K} = -85$, $E_\mathrm{Ca} = 120$, $E_\mathrm{H} = 20$.

The ion channel maximal conductances (expressed in mS/cm$^2$) are $\bar{g}_\mathrm{leak} = 0.055$, $\bar{g}_\mathrm{Na} = 170$, $\bar{g}_\mathrm{K,D} = 40$, $\bar{g}_\mathrm{Ca,T} = 0.55$, $\bar{g}_\mathrm{K,Ca} = 4$, $\bar{g}_\mathrm{H} = 0.01$.

The variables $m_\mathrm{ion}$ (resp. $h_\mathrm{ion}$) represents the activation (resp. inactivation) variable of the ion channel "ion". Their dynamics are given by

$$\tau_{m_\mathrm{ion}}(V)\frac{dm_\mathrm{ion}}{dt} = m_{\mathrm{ion},\infty}(V) - m_\mathrm{ion},$$
$$\tau_{h_\mathrm{ion}}(V)\frac{dh_\mathrm{ion}}{dt} = h_{\mathrm{ion},\infty}(V) - h_\mathrm{ion}.$$

The steady-state values $x_{\mathrm{ion},\infty}(V)$ and the time constants $\tau_{x_\mathrm{ion}}(V)$ of the ion channel "ion" are voltage-dependent such as:

$$x_{\mathrm{ion},\infty}(V) = \frac{1}{1 + \exp((V + V_{x_\mathrm{ion},\mathrm{half}})/\mathrm{slope}_{x_\mathrm{ion}})},$$
$$\tau_{x_\mathrm{ion}}(V) = A - \frac{B}{1 + \exp((V + V_{\tau_{x_\mathrm{ion}},\mathrm{half}})/\mathrm{slope}_{\tau_{x_\mathrm{ion}}})}.$$

The parameters for the different channels are given following the format $x_{\mathrm{ion},\infty}(V) = f(V_{x_\mathrm{ion},\mathrm{half}}, \mathrm{slope}_{x_\mathrm{ion}})$; $m_{\mathrm{Na},\infty} = f(35.5, -5.29)$, $h_{\mathrm{Na},\infty} = f(48.9, 5.18)$, $m_{\mathrm{K,D},\infty} = f(12.3, -11.8)$, $m_{\mathrm{Ca,T},\infty} = f(67.1, -7.2)$, $h_{\mathrm{Ca,T},\infty} = f(80.1, 5.5)$, $m_{\mathrm{H},\infty} = f(80.0, 6.0)$ and $\tau_{x_\mathrm{ion}}(V) = g(A, B, V_{\tau_{x_\mathrm{ion}},\mathrm{half}}, \mathrm{slope}_{\tau_{x_\mathrm{ion}}})$; $\tau_{m_\mathrm{Na}} = f(1.32, 1.26, 120, -25)$, $\tau_{h_\mathrm{Na}}(V) = (0.67/(1{+}\exp((V{+}62.9)/{-10.0})))\cdot(1.5{+}1/(1{+}\exp((V{+}34.9)/3.6)))$, $\tau_{m_\mathrm{K,D}} = g(0.2, 6.4, 28.3, -19.2)$, $\tau_{m_\mathrm{Ca,T}} = g(21.7, 21.3, 68.1, -20.5)$, $\tau_{h_\mathrm{Ca,T}} = g(410, 179.6, 55, -16.9)$, $\tau_{m_\mathrm{H}} = g(272., -1149., 42.2, -8.73)$.

In this conductance-based model, the calcium $\mathrm{Ca}^{2+}$ is entering through the T-type calcium channel. The dynamics of the calcium concentration is thus given by:

$$\frac{d[\mathrm{Ca}^{2+}]_\mathrm{T}}{dt} = -k_1 I_\mathrm{Ca,T} - k_2[\mathrm{Ca}^{2+}]_\mathrm{T},$$

where $k_1$ and $k_2$ are the rate variables. The kinetics for the synapse are $k_1 = 0.1$ M/msA, $k_2 = 0.01$ nM/ms. The calcium-activated potassium current considers this calcium entry to update its gating variable:

$$m_{\mathrm{K,Ca}}([\mathrm{Ca}^{2+}]_{\mathrm{T}}) = \left(\frac{[\mathrm{Ca}^{2+}]_{\mathrm{T}}}{[\mathrm{Ca}^{2+}]_{\mathrm{T}} + K_{\mathrm{D}}}\right)^2,$$

where $K_{\mathrm{D}}$ is a calcium-activation constant. The calcium half-activation constant is $K_{\mathrm{D}} = 170$nM.

**Network** We have three sets of neurons $\mathcal{N}_{\mathrm{inh}}$, $\mathcal{N}_{\mathrm{pre}}$, and $\mathcal{N}_{\mathrm{post}}$. The excitatory synaptic current perceived by the postsynaptic neuron $i$ from presynaptic neuron $j$ is characterized in the main methods. The inhibitory synaptic current perceived by the postsynaptic neuron $i$ from presynaptic neuron $j$ is characterized by

$$I_{\mathrm{GABA_A},ij} = \bar{g}_{\mathrm{GABA_A},ij} \cdot s_{\mathrm{GABA_A},j} \cdot (V_i - E_{\mathrm{GABA_A}}),$$
$$I_{\mathrm{GABA_B},ij} = \bar{g}_{\mathrm{GABA_B},ij} \cdot s_{\mathrm{GABA_B},j} \cdot (V_i - E_{\mathrm{GABA_B}}),$$

where $\bar{g}_{\mathrm{GABA_A},ij}$ and $\bar{g}_{\mathrm{GABA_B},ij}$ respectively represent the maximal conductance of the $\mathrm{GABA_A}$ receptors and the $\mathrm{GABA_B}$ receptors. Without variability, they are set to $2\,\mathrm{mS/cm^2}$ and $1.5\,\mathrm{mS/cm^2}$ . The variable $s_{\mathrm{GABA_A},j}$ denotes the gating variable of the $\mathrm{GABA_A}$ postsynaptic receptor ($\mathrm{GABA_A}$r), dynamically modulated by the presynaptic membrane voltage ($V_j$) and $E_{\mathrm{GABA_A}}$ is the reversal potential of $\mathrm{GABA_A}$r (set to $E_{\mathrm{Cl}} = -70\,\mathrm{mV}$)). The variable $s_{\mathrm{GABA_B},j}$ is the same for the the $\mathrm{GABA_B}$ postsynaptic receptor ($\mathrm{GABA_B}$r) where the reversal potential of $\mathrm{GABA_B}$r is set to $E_{\mathrm{K}} = -85\,\mathrm{mV}$.

The gating variables of the synapses are $s_{\mathrm{AMPA},j}$, $s_{\mathrm{GABA_A},j}$ and $s_{\mathrm{GABA_B},j}$ are variables whose dynamics depends on the considered presynaptic membrane potential following the equations

$$\dot{s}_{\mathrm{AMPA},j} = 1.1\,T_{\mathrm{m}}(V_j)(1 - s_{\mathrm{AMPA},j}) - 0.19\,s_{\mathrm{AMPA},j}$$
$$\dot{s}_{\mathrm{GABA_A},j} = 0.53\,T_{\mathrm{m}}(V_j)(1 - s_{\mathrm{GABA_A},j}) - 0.18\,s_{\mathrm{GABA_A},j}$$
$$\dot{s}_{\mathrm{GABA_B},j} = 0.016\,T_{\mathrm{m}}(V_j)(1 - s_{\mathrm{GABA_B},j}) - 0.0047\,s_{\mathrm{GABA_B},j}$$

with transmitter concentration follows the formalism described in (Destexhe et al., 1994), that is,

$$T_{\mathrm{m}}(V) = \frac{1}{1 + \exp\left[-(V - 2)/5\right]}.$$

The numerical values (1.1, 0.19, 0.53, 0.18, 0.016, and 0.0047) are rate parameters to mimic the kinetics of the synaptic receptors. These values originated from (Destexhe et al., 1994). The smallest, the slowest the kinetics.

#### Text S2 Plasticity implementation

In this section, the different plasticity rules and their variations are defined. The parameters are given in the code available on the GitHub https://github.com/KJacquerie/Burst-Attractor.

##### Calcium-based models

**Calcium dynamics** We consider the linear calcium dynamics suggested by (Graupner and Brunel, 2012; Graupner et al., 2016). The presynaptic and postsynaptic spike contributions add linearly. Indeed, calcium enters from NMDA receptors and voltage-dependent calcium channels. Instead of describing the whole calcium, the phenomenological effect on the calcium variation is considered. At each pre or postsynaptic spike (respectively named as $t_{j,k}$ and $t_{i,k}$), the calcium immediately rises and then exponentially decays characterized by a calcium decay time constant equal to $\tau_{\text{Ca}}$:

$$\frac{dc_j}{dt} = -\frac{c_j}{\tau_{\text{Ca}}} + C_{\text{pre}} \sum_{k \in \mathcal{T}_j} \delta(t - t_{j,k} - D),$$

$$\frac{dc_i}{dt} = -\frac{c_i}{\tau_{\text{Ca}}} + C_{\text{post}} \sum_{k \in \mathcal{T}_i} \delta(t - t_{i,k}),$$

where $C_{\text{pre}}$ and $C_{\text{post}}$ are the presynaptically and postsynaptically evoked calcium amplitudes. The parameter $D$ is a time-delay between the presynaptic spike and the corresponding postsynaptic calcium transient occurrence accounts for the slow rise time of the NMDAr-mediated calcium influx (Graupner and Brunel, 2012; Graupner et al., 2016).

The total calcium amplitude $c_{ij}(t)$ driving the synaptic change is given by:

$$c_{ij}(t) = c_j(t) + c_i(t).$$

The time-evolution for several pre- and postsynaptic spiking activity is written such as (Graupner et al., 2016):

$$c_{ij}(t) = \sum_{k \in \mathcal{T}_j} C_{\text{pre}} \exp\left(\frac{t - t_{j,k} - D}{\tau_{\text{Ca}}}\right) + \sum_{k \in \mathcal{T}_i} C_{\text{post}} \exp\left(\frac{t - t_{i,k}}{\tau_{\text{Ca}}}\right).$$

The resting calcium concentration is set to zero. The calcium concentrations are dimensionless. Both simplification is acknowledged because the synaptic rules are adapted in accordance. If a resting calcium concentration is wanted, the thresholds of potentiation and depression will be adapted. This notation follows the original paper notation.

**[Graupner et al. 2012]** Graupner and Brunel developed a breakthrough calcium-dependent synaptic rule. It implements two opposing calcium-triggered pathways leading to an increase or a decrease in synaptic strength. Indeed, potentiation and depression are activated above calcium thresholds. It is defined by Equation 3

The parameters are fitted for the frequency plasticity-induced protocol (Sjöström et al., 2001) in the cortex (CTX) or the spike-time dependent plasticity-induced protocol (Bi and Poo, 1998) in the hippocampus (HPC).

**[Graupner et al. 2016]** This model evolves in (Graupner et al., 2016) such as the synaptic plasticity rule becomes Equation 4

The parameters are fitted on CTX data in (Graupner et al., 2016). We have also matched the plasticity parameters to reproduce the data obtained in HPC.

This synaptic rule can be easily written using a standard form for ordinary differential equations with a steady-state value and a time constant for each calcium region as shown in Equation 5

**[Shouval et al. 2012]** In (Shouval et al., 2002), the synaptic change follows a typical first-order differential equation:

$$\dot{w}_{ij} = \frac{1}{\tau_w(c_{ij})}\left(\Omega(c_{ij}) - w_{ij}\right),$$

where the time constant $\tau_w(c_{ij})$ and the steady-state value $\Omega(c_{ij})$ are calcium-dependent. This steady state value is defined by two sigmoids in order to build the U-shape for $\Omega$:

$$\Omega(c_{ij}) = a_0 - a_0 \exp\left(\frac{b_1(c_{ij}-a_1)}{1+\exp(b_1(c_{ij}-a_1))}\right) + m_2 \exp\left(\frac{b_2(c_{ij}-a_2)}{1+\exp(b_2(c_{ij}-a_2))}\right).$$

This expression relies on five parameters:

- $a_0$ is the ordinate at low levels of calcium;
- $a_1$ is the abscissa where the ordinate $a_0$ is divided by 2 (it dictates the place along the x-axis where the first sigmoid is decreasing);
- $b_1$ governs the sharpness of the decrease around $a_1$ (the bigger, the flatter the slope);
- $m_2$ the converging value at high calcium level;
- $a_2$ is the x-value where $\Omega$ equal $m_2/2$;
- $b_2$ is similar as $b_1$ (it dictates the sharpness of the slope).

The time constant is also calcium-dependent and it is given by (Shouval et al., 2002)

$$\tau_w(c_{ij}) = P_4 + \frac{P_1}{P_2 + c_{ij}^{P_3}},$$

where $P_1$, $P_2$, $P_3$, and $P_4$ are fitted parameters to describe the calcium-dependent time constant. The different parameters were identified to reproduce the different potentiation and depression levels provided by the previous model for the frequency-dependency protocol (CTX, soft bounds). The synaptic rule is using the calcium dynamics presented above. The parameters are $a_0 = 0.5$, $a_1 = 1.31$, $a_2 = 1.8$, $b_1 = 20$, $b_2 = 40$, $P_1 = 4e3$, $P_2 = P1.1e-6$, $P_3 = 2.4$, $P_4 = 1$. For hard bounds, the parameters become $m_1 = 0.25$, $a_1 = 1$, $a_2 = 2$, $b_1 = 40$, $b_2 = 10$, $m_2$, $P_1, P_2, P_3, P_4$ do not change.

**[Deperrois et al. 2020]** The synaptic weight influences the postsynaptic calcium dynamics as mentioned in (Deperrois and Graupner, 2020). It is translated by a linear scale coupling between the presynaptically induced calcium amplitude:

$$\frac{dc_j}{dt} = -\frac{c_j}{\tau_{\text{Ca}}} + w_{ij}C_{\text{pre}}\sum_k \delta(t - t_{j,k} - D).$$

The effect of the short-term depression (STD) at the presynaptic site as described in (Deperrois and Graupner, 2020) is also tested in our project. To account for the available presynaptic resources, (Deperrois and Graupner, 2020) uses a variable $x$. Then, $U$ considers the fraction of the resources required at each presynaptic spike and $\tau_{\text{rec}}$ is the resource recovery time constant for the variable $x$ to come back to its resting state equal to 1

$$\frac{dx}{dt} = \frac{1-x}{\tau_{\text{rec}}} - Ux\sum_{k\in\mathcal{T}_j} \delta(t - t_{j,k} - D).$$

So we have

$$\frac{dc_j}{dt} = -\frac{c_j}{\tau_{\text{Ca}}} + w_{ij}C_{\text{pre}}Ux\sum_{k\in\mathcal{T}_j} \delta(t - t_{j,k} - D).$$

The parameters are fitted for the frequency-dependent plasticity-induced protocol (CTX) in soft or hard bounds, without short-term depression $\tau_{Ca} = 32.19$, $C_{\text{pre}} = 1.61$, $C_{\text{post}} = 1.124$, $D = 5.7527$, $\tau_w = 79\,975\,\text{ms}$, $\theta_{\text{p}} = 1.63$, $\theta_{\text{d}} = 1$, $\gamma_{\text{p}} = 161.99$, $\gamma_{\text{d}} = 31.976$; with short-term depression $\tau_{Ca} = 38.35$, $C_{\text{pre}} = 3.99$, $C_{\text{post}} = 1.29$, $D = 9.24$, $\tau_w = 299\,877.8\,\text{ms}$, $\theta_{\text{p}} = 1.63$, $\theta_{\text{d}} = 1$, $\gamma_{\text{p}} = 564.4$, $\gamma_{\text{d}} = 111.3$, $\tau_{\text{rec}} = 148.92$, and $U = 0.3838$.

**Hard-bound implementation** As pioneered in (Shouval et al., 2002), calcium drives the synaptic change. The first implementation suggested was:

$$\dot{w}_{ij} = \rho\ \Omega(c_{ij}).$$

The synaptic change follows the speed given by $\Omega$. It leads to weight runaway. The simplest solution to overcome this runaway is the addition of "hard bound" to constrain the synaptic weight between lower and an upper limit. In this work, they are respectively fixed at 0 and 1. Computationally, it is implemented such as: $\text{if}(w_{ij} \geq 1) : w_{ij} = 1; \text{if}(w_{ij} \leq 0) : w_{ij} = 0$.

We transformed the two-thresholds model suggested by (Graupner et al., 2016). The potentiation and depression terms ($\gamma_\text{p}$ and $\gamma_\text{d}$) are dependent on the synaptic weights. In phenomenological models, it is called 'soft bounds'. A strong weight has a weaker effective potentiation rate and a stronger effective depression rate. By contrast, a weak weight has a stronger effective potentiation rate and a weaker depression rate. Mathematically, it is easily observed from the equation: $\tau_w \dot{w}_{ij} = \gamma_\text{p}(1 - w_{ij}) - \gamma_\text{d} w_{ij}$. For a strong weight equal to 0.9: the expression becomes: $\tau_w \dot{w}_{ij} = \gamma_\text{p}0.1 - \gamma_\text{d}0.9$. The effective potentiation rate is 10% the original value while the effective depression rate is 90 % is the original value.

To overcome this weight dependency, we convert the soft bounds expression into hard bounds:

$$\begin{cases} \tau_w \dot{w}_{ij} = \Omega_0, & \text{if } c_{ij} < \theta_\text{d}, \\ \tau_w \dot{w}_{ij} = -\Omega_\text{d} = -0.5\gamma_\text{d}, & \text{if } \theta_\text{d} \leq c_{ij} \leq \theta_\text{p}, \\ \tau_{w,\text{p}} \dot{w}_{ij} = \Omega_\text{p} = 0.5(\gamma_\text{p} - \gamma_\text{d}), & \text{if } \theta_\text{p} \leq c_{ij}. \end{cases}$$

At low levels of calcium, the synaptic weight is unchanged ($\Omega_0 = 0$). At intermediate levels, we used the expression provided in the original model by considering a fixed mean weight equal to 0.5. Therefore, the synaptic change is decreased by a depression rate of $\frac{0.5\gamma_\text{d}}{\tau_w}$. At high levels of calcium, the potentiation rate equals $\frac{0.5(\gamma_\text{p} - \gamma_\text{d})}{\tau_w}$. This transformation is valid because the provided potentiation rate is bigger than the depression rate (parameters originate from the paper). The presence of 0.5 comes from the removal of $w$-dependency in the main equation. To do so, its medium value has been validated during the fitting protocol in experimental data (Sjöström et al., 2001). Keeping $\gamma_\text{d}$ for the depression rate and $\gamma_\text{p}$ for the potentiation rate leads to a discrepancy with the experimental data mentioned.

The same development is performed for (Shouval et al., 2002). The synaptic rule simply becomes: $\tau_w(c_{ij})\dot{w}_{ij} = \Omega(c_{ij})$ with the addition of hard bounds. Similar mathematical manipulations have been done in calcium-based rules established by considering the presynaptic resource depletion. The weight dependency has been removed inside the synaptic rule and the parameters have been updated. The parameters were once again fitted to reproduce the frequency-dependency protocol from (Sjöström et al., 2001).

##### Spike-time dependent models

**Pair-based model** To reproduce the classical STDP window, pair-based model is implemented using synaptic traces, respectively $x$ and $y$ for the presynaptic neuron $j$ and postsynaptic neuron $i$:

$$\frac{dx_{ij}}{dt} = -\frac{x_{ij}}{\tau_x} + \delta\left(t - t_i\right)$$
$$\frac{dy_{ij}}{dt} = -\frac{y_{ij}}{\tau_y} + \delta\left(t - t_j\right).$$

The weight change relative to the STDP window was then computed using the following equations, with explicit hard bounds defined such as $0 < w_{ij} < 1$:

$$w_{ij}(t) \rightarrow \begin{cases} w_{ij}(t) + A^+ x_{ij}(t), & \text{at } t = t_j, \\ w_{ij}(t) - A^- y_{ij}(t), & \text{at } t = t_i, \end{cases}$$

or using soft bounds:

$$w_{ij}(t) \rightarrow \begin{cases} w_{ij}(t) + A^{+}(1 - w_{ij})x_{ij}(t), & \text{at } t = t_j, \\ w_{ij}(t) - A^{-}w_{ij}y_{ij}(t), & \text{at } t = t_i, \end{cases}$$

where $A^{+} > 0$, $A^{-} > 0$ and $t_i$ (resp. $t_j$) representing the spike timing of the presynaptic (resp. postsynaptic) neuron, *i.e.*, each time $t = t_i$ (resp. $t = t_j$), we consider a post-pre (resp. pre-post) pair for the weight change (Morrison et al., 2008; Song et al., 2000; Van Rossum et al., 2000). In this project, we combine soft-bound and hard-bound dependency using the notation proposed in (Gütig et al., 2003):

$$w_{ij} \rightarrow \begin{cases} w_{ij} + A^{+}(1 - w_{ij})^{\mu} e^{-\Delta t/\tau^{+}}, & \text{at } t_i \text{ if } t_j < t_i, \\ w_{ij} - A^{-}w_{ij}^{\mu} e^{\Delta t/\tau^{-}}, & \text{at } t_j \text{ if } t_j > t_i, \end{cases}$$

where $\mu$ is equal to 0 for hard-bounds and 1 for soft-bounds.

The parameters are $A^{+}$=0.0096, $A^{-}$=0.0053, $\tau_x$= 16.8 and $\tau_y$=33.7 (HPC) (Bi and Poo, 2001).

**Triplet model** Similarly to the pair-based model, the triplet model was implemented using trace variables following (Pfister and Gerstner, 2006):

$$\begin{aligned} \frac{dx_1}{dt} &= -\frac{x_1}{\tau^{+}} + \delta(t - t_i) \\ \frac{dx_2}{dt} &= -\frac{x_2}{\tau_x} + \delta(t - t_i) \\ \frac{dy_1}{dt} &= -\frac{y_1}{\tau^{-}} + \delta(t - t_j) \\ \frac{dy_2}{dt} &= -\frac{y_2}{\tau_y} + \delta(t - t_j), \end{aligned}$$

where $t_i$ (resp. $t_j$) is the timing of a presynaptic spike (resp. postsynaptic). The full model implemented by (Pfister and Gerstner, 2006) takes into account weight change due to pre-post, with the constant $A_2^{+}$, inducing potentiation or post-pre pairs, with the constant $A_2^{-}$, inducing depression (similar to classical pair-based model, with $x_1(t)$ and $r_2(t)$ as the presynaptic and postsynaptic traces, respectively with their respective time constant $\tau^{+}$ and $\tau^{-}$).

The improvement over the classic pair-based model is that a triplet of spikes is also considered. Thanks to previously introduced traces $y_2$, decaying with a time constant $\tau_y$, and $x_2$, decaying with $\tau_x$, pre-post-pre triplets are treated (associated with the constant $A_3^{-}$, inducing depression) as well as post-pre-post triplets (associated with the constant $A_3^{+}$, inducing potentiation).

$$w_{ij}(t) \rightarrow \begin{cases} w_{ij}(t) + x_1(t)\left[A_2^{+} + A_3^{+}o_2(t - \epsilon)\right], & \text{at } t = t_j, \\ w_{ij}(t) - o_1(t)\left[A_2^{-} + A_3^{-}x_2(t - \epsilon)\right], & \text{at } t = t_i. \end{cases}$$

The parameters are for the minimal model (CTX) $A_2^{+} = 0$, $A_3^{+} = 6.5e^{-3}$, $A_2^{-} = 7.1e^{-3}$, $A_3^{-} = 0$, $\tau_x = 101\,\text{ms}$, $\tau_y = 125\,\text{ms}$, $\tau^{+} = 16.8\,\text{ms}$, $\tau^{-} = 33.7\,\text{ms}$ and (HPC) $A_2^{+} = 5.3e^{-3}$, $A_3^{+} = 8e^{-3}$, $A_2^{-} = 3.5e^{-3}$, $A_3^{-} = 0$, $\tau_x = 101\,\text{ms}$, $\tau_y = 40\,\text{ms}$, $\tau^{+} = 16.8\,\text{ms}$, $\tau^{-} = 33.7\,\text{ms}$.

The model can be described using soft bounds (Graupner et al., 2016):

$$w_{ij}(t) \rightarrow \begin{cases} w_{ij}(t) + x_1(t)(1 - w_{ij})\left[A_2^{+} + A_3^{+}o_2(t - \epsilon)\right], & \text{at } t = t_j, \\ w_{ij}(t) - o_1(t)w_{ij}\left[A_2^{-} + A_3^{-}x_2(t - \epsilon)\right], & \text{at } t = t_i. \end{cases}$$

The parameters are (CTX) $A_2^{+} = 0$, $A_3^{+} = 0.0165746$, $A_2^{-} = 0.00826477$, $A_3^{-} = 0$, $\tau_x = 56.38\,\text{ms}$, $\tau_x = 101\,\text{ms}$, $\tau_y = 56.3824\,\text{ms}$, same $\tau^{+}$, $\tau^{-}$ and for (HPC) same as for hard-bounds.

### Text S3 Computational experiments: numerical values

The parameters associated with current and initial synaptic weights are constant in the different simulations: $I_{\text{app,inh}}(\text{Tonic}) = 3\,\text{nA/cm}^2$, $I_{\text{app,inh}}$ (Burst) $= -1.2\,\text{nA/cm}^2$, $w_0$=0.5, $\bar{g}_{\text{AMPA}}$=0.001.

The parameters associated with the synaptic plasticity are given in (Graupner and Brunel, 2012) fitting cortical (CTX) data (Sjöström et al., 2001): $\tau_{\text{Ca}} = 22.6936\,\text{ms}$, $C_{\text{pre}} = 0.56$, $C_{\text{post}} = 1.24$, $D = 4.60\,\text{ms}$, $\tau_w = 346.3615 \times 10^3\,\text{ms}$, $\gamma_{\text{p}} = 725.085 \times 1.1$ (Tonic), $\gamma_{\text{p}} = 725.085 \times 0.95$ (Burst), $\gamma_{\text{d}} = 331.909$, $\theta_{\text{p}} = 1.3$, $\theta_{\text{d}} = 1$, $w^* = 0.5$. The potentiation rate $\gamma_p$ is slightly scaled up during tonic firing to induce stronger potentiation compared to the initial model, and it is reduced by 5 % during burst firing to place the fixed-point at a lower value compared to the initial model.

For (Graupner and Brunel, 2012) fitting hippocampus (HPC) data (Bi and Poo, 1998): $\tau_{\text{Ca}} = 20\,\text{ms}$, $C_{\text{pre}} = 1$, $C_{\text{post}} = 2$, $D = 13.7\,\text{ms}$, $\tau_w = 150 \times 10^3\,\text{ms}$, $\gamma_{\text{p}} = 321.808$ $\gamma_{\text{d}} = 200$, $\theta_{\text{p}} = 1.3$, $\theta_{\text{d}} = 1$, $w^* = 0.5$.

The parameters associated with the synaptic plasticity are given in (Graupner et al., 2016) fitting cortical (CTX) data (Sjöström et al., 2001): $\tau_{\text{Ca}} = 22.272\,12\,\text{ms}$, $C_{\text{pre}} = 0.8441$, $C_{\text{post}} = 1.62138$, $D = 9.537\,09\,\text{ms}$, $\tau_w = 520\,761.29\,\text{ms}$, $\gamma_{\text{p}} = 597.08922$, $\gamma_{\text{d}} = 137.7586$, $\theta_{\text{p}} = 2.009289$, $\theta_{\text{d}} = 1$. For (Graupner et al., 2016) in soft bounds that fit the hippocampal data of (Bi and Poo, 1998), the parameters remain the same, except that $\theta_p$ becomes equal to 1.45.

The parameters used in each simulation are for Fig 1 $N = 50, M = 50, T_{\text{state}} = 20\,\text{s}, N_{\text{state}} = 8$, for Fig 2 $N = 50, M = 50, T_{\text{state}} = 50\,\text{s}$, for Fig 3 $N = 50, M = 50$, $T_{\text{state}} = 20\,\text{s}$, and for Fig 4 $N_{\text{state}} = 4$, $N = 1$, $M = 1$,$T_{\text{state}} = 20\,\text{s}$ .

### Text S4 Derivation of the burst-induced attractor in a calcium-based model

We consider the calcium-based plasticity model of Graupner and Brunel (Graupner et al., 2016), given in Equation 4. This rule specifies how the synaptic weight $w_{ij}$ evolves as a function of the postsynaptic calcium concentration $c_{ij}$. Depending on the calcium level, the dynamics are governed by three regimes: when calcium remains below the depression threshold ($c_{ij} < \theta_{\text{d}}$), the weight does not change; at intermediate calcium levels ($\theta_{\text{d}} \leq c_{ij} < \theta_{\text{p}}$), the weight relaxes toward the depression steady state $\Omega_{\text{d}} = 0$ with time constant $\tau_{w,\text{d}} = \tau_w/\gamma_{\text{d}}$; and at high calcium concentrations ($c_{ij} \geq \theta_{\text{p}}$), the weight relaxes toward the potentiation steady state $\Omega_{\text{p}} = \gamma_{\text{p}}/(\gamma_{\text{p}}+\gamma_{\text{d}})$ with time constant $\tau_{w,\text{p}} = \tau_w/(\gamma_{\text{p}}+\gamma_{\text{d}})$. These regimes are summarized in:

$$\begin{cases} \dot{w}_{ij} = 0, & \text{if } c_{ij} < \theta_{\text{d}}, \\ \tau_{w,\text{d}}\dot{w}_{ij} = \Omega_{\text{d}} - w_{ij}, & \text{if } \theta_{\text{d}} \leq c_{ij} < \theta_{\text{p}}, \\ \tau_{w,\text{p}}\dot{w}_{ij} = \Omega_{\text{p}} - w_{ij}, & \text{if } \theta_{\text{p}} \leq c_{ij}. \end{cases} \tag{5}$$

Because calcium fluctuates on the timescale of individual spikes and bursts (milliseconds), while synaptic weights evolve on much slower timescales (seconds to minutes), we reformulate the dynamics by averaging over an intermediate window of length $T$. The interval $T$ must be long enough to capture typical calcium fluctuations. The averaged dynamics are then given by

$$\dot{w}_{ij} = \frac{1}{T}\int_t^{t+T} \frac{1}{\tau_w(c_{ij}(s))}\left[\Omega(c_{ij}(s)) - w_{ij}\right] ds,$$

where $\Omega(c_{ij})$ denotes the steady-state value (either $\Omega_{\text{d}}$ or $\Omega_{\text{p}}$) associated with a given calcium regime, and $\tau_w(c_{ij})$ is the corresponding time constant. This formulation describes the effective pull exerted on the synaptic weight by the sequence of calcium transients encountered during the interval.

To express this average more explicitly, we define the effective times spent in each regime. The effective depression time $\alpha_{\text{d},ij}$ is the fraction of the interval spent between the depression and potentiation thresholds, normalized by $\tau_{\text{d}}$. Similarly, the effective potentiation time $\alpha_{\text{p},ij}$ measures the fraction of time spent above the potentiation threshold, normalized by $\tau_{\mathrm{p}}$:

$$\alpha_{\mathrm{d},ij} = \frac{1}{\tau_{\mathrm{d}}}\frac{1}{T}\int_t^{t+T} \Theta(\theta_{\mathrm{p}} - c_{ij}(s))\,\Theta(c_{ij}(s) - \theta_{\mathrm{d}})\,ds, \qquad \alpha_{\mathrm{p},ij} = \frac{1}{\tau_{\mathrm{p}}}\frac{1}{T}\int_t^{t+T} \Theta(c_{ij}(s) - \theta_{\mathrm{p}})\,ds,$$

where $\Theta$ is the Heaviside step function. Periods where calcium remains below $\theta_{\mathrm{d}}$ not contribute to a change and are quantified by $\alpha_0$. Fig 2A illustrates how these contributions are measured for a representative calcium trace.

With these definitions, the averaged dynamics is simplified to

$$\dot{w}_{ij} = \alpha_{\mathrm{d},ij}(\Omega_{\mathrm{d}} - w_{ij}) + \alpha_{\mathrm{p},ij}(\Omega_{\mathrm{p}} - w_{ij}),$$

which makes explicit that the weight is simultaneously pulled toward the depression and potentiation steady states, with strengths proportional to the effective times spent in each regime. The fixed point $\bar{w}_{ij}$ is reached when these contributions balance, yielding Equation 1. This expression shows that the long-term value of the synaptic weight is a weighted average of the potentiation and depression steady states, where the weights reflect how often calcium crosses each threshold (Gütig et al., 2003; Dorman and Blackwell, 2021). In other words, the precise statistics of calcium fluctuations determine the effective balance of potentiation and depression, and thus the fixed point to which the synapse converges.

#### Text S5 Derivation of the burst-induced attractor in a spike-time dependent plasticity rule

We next derive the fixed point for a spike-timing–dependent plasticity (STDP) rule. The pair-based model considers presynaptic and postsynaptic spike times (resp. $t_i$ and $t_j$), with their relative timing with a time difference $\Delta t = t_j - t_i$ to induce the change in synaptic weight. A classical pair-based STDP window is used: when a presynaptic spike precedes a postsynaptic spike, the weight increases (potentiation), and when the order is reversed, the weight decreases (depression) (Abbott and Nelson, 2000; Song et al., 2000; Morrison et al., 2008; Rubin et al., 2001; Van Rossum et al., 2000):

$$w \to \begin{cases} w_{ij} + A^{+}(1 - w_{ij})^{\mu} e^{-\Delta t/\tau^{+}}, & \text{at } t_j \text{ if } t_i < t_j, \\ w_{ij} - A^{-} w^{\mu} e^{\Delta t/\tau^{-}}, & \text{at } t_i \text{ if } t_i > t_j, \end{cases}$$

where $A^{+}$ and $A^{-}$ are the potentiation and depression parameters, $\mu$ stands for the weight-dependency, $e^{-|s|/\tau^{\pm}}$ stands for the STDP kernel in potentiation or depression with $\tau^{+}$ and $\tau^{-}$ being the time constants given in the pair-based model. The plasticity parameters $A^{+} > 0$, $A^{-} > 0$ (Morrison et al., 2008; Song et al., 2000). The weight dynamics can be constrained in two manners; either by using *hard bounds* or *soft bounds*. Hard bounds permit to stop the weight increase or decrease by adding upper or lower limits. Soft bounds decelerate the evolution if the weight reaches a bound. It is modeled by the weight-dependency parameter $\mu$: $\mu$ is equal to 0 for hard-bounds and to 1 for soft-bounds) (Gerstner and Kistler, 2002).

The functions $e^{-|\Delta t|/\tau^{\pm}}$ are the temporal kernel of potentiation and depression. If we introduce $S_i(t) = \sum_k \delta(t - t_{i,k})$ and $S_j(t) = \sum_k \delta(t - t_{j,k})$ for the spike trains of presynaptic neuron $j$ and the postsynaptic neuron $i$, the evolution of the synaptic weight can be written as follows:

$$\dot{w}_{ij} = -A^{-} w_{ij}^{\mu} \left[\int_{-\infty}^{0} e^{s/\tau^{-}} S_j(t-s)ds\right] S_i(t) + A^{+}(1 - w_{ij})^{\mu} \left[\int_{0}^{\infty} e^{-s/\tau^{+}} S_i(t-s)ds\right] S_j(t). \quad (6)$$

The time evolution of a weight and its convergence occurs on a time interval much larger than typical interspike intervals. Therefore, following the work of (Gütig et al., 2003; Legenstein and Maass, 2005), we can average the dynamics of the synaptic weight over a time interval $T$ and get

$$\dot{w}_{ij} = -A^{-} w_{ij}^{\mu} \int_{-\infty}^{0} e^{s/\tau^{-}} C(s;t)ds + A^{+}(1 - w_{ij})^{\mu} \int_{0}^{\infty} e^{-s/\tau^{+}} C(s;t)ds,$$

where $C(s;t)$ is the (temporally averaged) correlation function between the pre and post spike trains, respectively noted $S_i(t) = \sum_k \delta(t - t_{i,k})$ and $S_j(t) = \sum_k \delta(t - t_{j,k})$, that is,

$$\lambda_{ij} = \alpha_\mathrm{p} \Omega_\mathrm{p} + \alpha - \mathrm{d}\Omega_\mathrm{d}.$$

For the spike-time dependent plasticity model (Song et al., 2000), the slope can be predicted by the equation

$$\lambda = A^+ C^+ - A^- C^-.$$

### Supplementary Figures

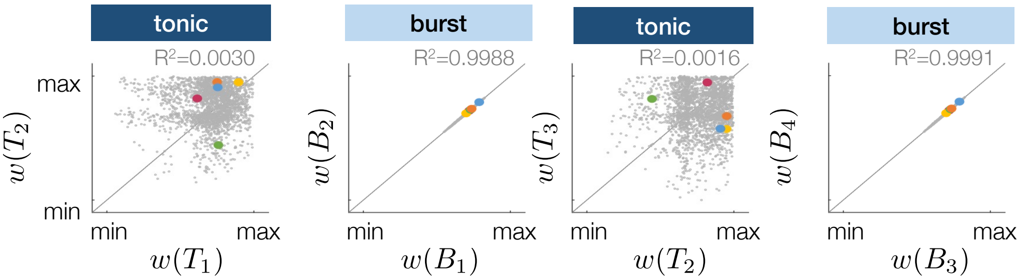


**Fig. S1:** Replication of the analysis illustrated in Fig 1D-E at different tonic and burst states.

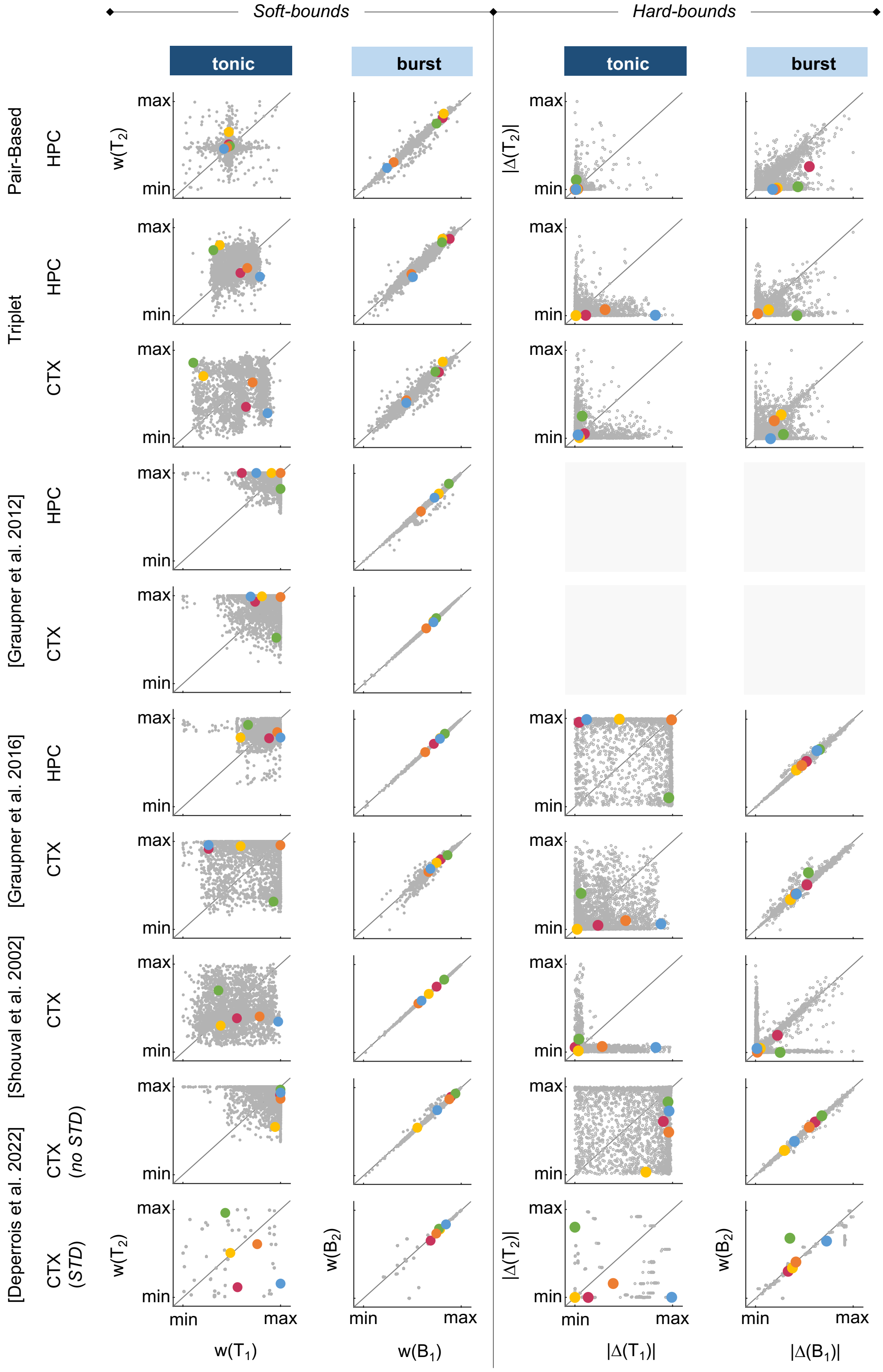


**Fig. S2:** Replication of the experiment illustrated in Fig 1C in various synaptic plasticity rules using soft-bounds (left panel) and hard-bounds (right panel). Comparison of the synaptic weights at the end of the third and fourth tonic firing states (left column) or the third and fourth burst firing states (right column), normalized between the minimal and maximal values. (CTX=cortex, data fitted on (Sjöström et al., 2001); HPC=hippocampus, data fitted on (Bi and Poo, 1998))

Fig S3 shows the evolution of synaptic weights between two excitatory neurons during burst firing for different initial conditions (0:0.1:1). In blue, trajectories correspond to unmodulated plasticity parameters, while in yellow they show neuromodulated parameters. The color gradient emphasizes the initial strength of the synaptic weights, with darker shades indicating larger initial values.

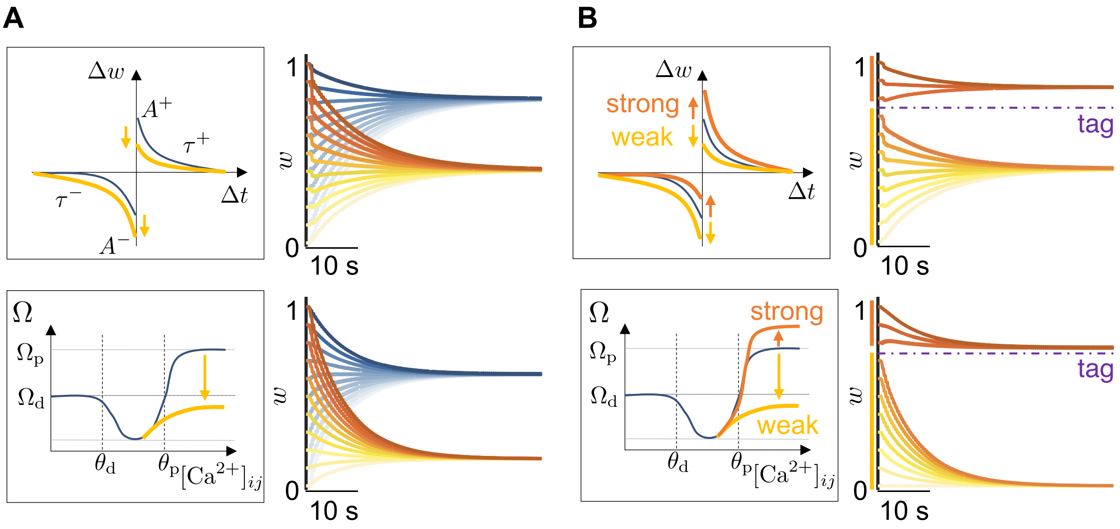


**Fig. S3:** A. Global neuromodulation. Left: plasticity rules in spike-based (top) and calcium-based (bottom) formulations, where potentiation and depression parameters ($A^p$,$A^m$ , or $\Omega_\mathrm{p}$, $\Omega_\mathrm{d}$) are globally downscaled (yellow arrows). Right: synaptic weights $w$ converge toward a lower attractor during bursting, showing that global parameter changes shift the attractor in weight space. B. Tag-dependent modulation. Left: plasticity rules where synapses above a tagging threshold receive enhanced potentiation parameters, while weaker synapses are downscaled. Right: tagged synapses converge to a higher attractor (purple dashed line), while untagged synapses decay toward a lower one, resulting in bimodal consolidation. This mechanism shows how neuromodulation and tagging can selectively stabilize strong synapses while weakening weaker ones.
